# Beyond Letters: Optimal Transport as a Model for Sub-Letter Orthographic Processing

**DOI:** 10.1101/2024.11.11.622929

**Authors:** Jack E. Taylor, Rasmus Sinn, Cosimo Iaia, Christian J. Fiebach

## Abstract

Letter processing plays a key role in visual word recognition. However, word recognition models typically overlook or greatly simplify early perceptual processes of letter recognition. We suggest that optimal transport theory may provide a computational framework for describing letter shape processing. We use representational similarity analysis to show that optimal transport cost (Wasserstein distance) between pairs of letters aligns with neural activity elicited by visually presented letters <225 ms after stimulus onset, outperforming an existing approach based on shape overlap. We additionally show that optimal transport can capture the emergence of geometric invariances (e.g., to position or size) observed in letter perception. Finally, we demonstrate that Wasserstein distance predicts neural activity similarly well to features from artificial networks trained to classify images and letters. However, whereas representations in artificial neural networks emerge in a computationally unconstrained manner, our proposal provides a computationally explicit route to modeling the earliest orthographic processes.

---

Reading is one of the most important cultural inventions humans have developed, for which the majority of humans rely on visual perception and recognition of written words. Visual word recognition processes allow visually presented words to be mapped to corresponding phonological, lexical, and semantic representations. To enable this, our brains must first extract and process orthographic information, identifying, recognising, and combining strokes, letters, and word forms in visual input. It is well known that such processes are guided by higher-level orthographic knowledge. For instance, letters are more easily recognised in the context of real words than they are in pseudowords or nonwords (Cattell, 1886b; Lally & Rastle, 2023; Reicher, 1969; Wheeler, 1970), and, when words are presented only briefly or at small visual angles, observers incorrectly report the presence of letters which are expected but are in fact absent (Jordan et al., 1999; Jordan et al., 2015; Pillsbury, 1897). Nevertheless, it is clear that bottom-up processing of component letters plays a substantial role in the recognition of visual words (Brossette et al., 2022; Grainger, 2008; Lally & Rastle, 2023; Pelli et al., 2003; Rosa et al., 2016). Indeed, neuroimaging evidence suggests that sensitivities to abstract letter identities, letter combinations, and then word forms peak successively over time, as information is passed on from more posterior, hierarchically ‘earlier’, to more anterior brain regions, along the ventral visual stream (Agrawal & Dehaene, 2025; Gutiérrez-Sigut et al., 2019; Lochy et al., 2018; Petit et al., 2006; Thesen et al., 2012; Vinckier et al., 2007; Woolnough et al., 2021).

In alphabetic scripts, letters thus provide a vital bridge between vision and language, and letter-shape processing can be conceptualised as the primary stage of orthographic processing. Despite this, the processes that comprise letter recognition have been largely overlooked in cognitive models of word recognition processes (Finkbeiner & Coltheart, 2009). When investigated experimentally, orthographic processes in word recognition are typically captured by orthographic measures whose descriptions begin at the level of letter identities (or, graphemes). For instance, the effect of the regularity of a word’s spelling on lexical decision performance, wherein more regular words are more easily recognised as real words, is captured by measures of orthographic neighbourhood density (Coltheart et al., 1977; Yarkoni et al., 2008), which describe how similar a given word’s string of letters is to those of other words in the same language or script. However, these measures limit themselves to treating letters as functionally irreducible categories, neglecting sub-graphemic (i.e., *graphetic;* Meletis & Dürscheid, 2022; Rezec, 2013) information. For instance, Coltheart’s *N* (Coltheart et al., 1977) quantifies a given word’s orthographic neighbourhood density as the number of other words of equal length that share all but one letter. This implies that orthographic similarity between any two words is binary; they are either neighbours at a distance of one letter substitution, or not, irrespective of letter identities. Similarly, the generalisation of this approach provided by orthographic Levenshtein distance (Yarkoni et al., 2008) counts the minimum number of letter insertions, deletions, and substitutions required to convert one word into another. Such approaches overlook entirely the graphetic features from which letters are composed, and the contribution that (dis-)similarities among these low-level perceptual and orthographic features may make to word recognition processes.

Graphetic processing has also been greatly simplified, or overlooked entirely, in computational models of word recognition. As with measures of orthographic neighbourhood density, such models instead typically begin their descriptions at the level of letters’ identities and their ordinal configurations within words (Norris, 2013; Snell, 2024). In one early and influential exception, Rumelhart and Siple (1974) pioneered an approach to modeling the processing of letters’ composite strokes with an artificial monospace type font comprised of just 14 character segments of fixed position and dimensions. Via combinations of these 14 segments, all 26 Latin alphabetic characters can be approximated. The Rumelhart-Siple font enabled them to incorporate stroke-level information into an interactive activation model of visual word recognition, implemented via an artificial neural network (ANN), without having to account for the shapes of real-world fonts. This innovation has remained a cornerstone in computational models of visual word recognition (e.g., Coltheart et al., 2001; Davis, 2010; Heilbron et al., 2020; McClelland & Rumelhart, 1981; Perry et al., 2007) that descend from the connectionist, interactive activation approach, and provides a framework that is extensible, supporting, for instance, descriptions of non-alphabetic scripts (Perfetti et al., 2005). And yet, models using variants of the Rumelhart-Siple font are, by definition, artificial, imposing synthetic structure and uniformity onto the diverse and heterogeneous typography of real-world letters. This limitation constrains the extent to which such models, and the orthographic similarities that can be extracted from them, reflect fully the true computations involved in orthographic processing.

More recent connectionist approaches include training ANNs to recognise real-world letters and words from raster-image representations (Agrawal & Dehaene, 2024; Chang et al., 2012; Hannagan et al., 2021). Rather than imposing artificial constraints on the input, graphetic representations in such ANNs are *emergent*, contingent on the model’s architecture, training regime, training sets, and hyperparameters. In one recent and notable study, Agrawal and Dehaene (2024) trained CORnet-Z, a convolutional neural network designed to reflect aspects of the primate ventral visual stream (Kubilius et al., 2018), on images of first, objects, and then, words. This study convincingly reproduced key effects observed in the so-called visual word form area, a region of the ventral occipitotemporal cortex (vOT) that has been consistently implicated in visual word recognition and orthographic processes (Dehaene & Cohen, 2011; Price, 2012; Woolnough et al., 2021). Agrawal and Dehaene showed that training the CORnet-Z architecture reproduced, in the late layers of the model thought to best capture particularly high-level ventral visual stream processing, the sensitivity to visual words observed in vOT, including its invariance to letter case, font, position, and size. Agrawal and Dehaene further compared the model’s representations to those of literate humans via a representational similarity analysis (RSA; Kriegeskorte et al., 2008) approach. It was shown that the more low-level, retinotopic representations, in the CORnet-Z layer taken to reflect the primary visual cortex, predicted magnetoencephalographic (MEG) event-related fields elicited by strings of letters for a sustained period, from 60 ms until at least 500 ms post stimulus onset (although the infinite-impulse-response 1 Hz high-pass filter applied in preprocessing is likely to have distorted the component’s onset somewhat; Rousselet, 2012; Tanner et al., 2015; VanRullen, 2011). A later layer in the model, taken to reflect subsequent processing in the inferotemporal cortex, predicted brain activity for a later sustained period, beginning at 140 ms post-stimulus, although partialling out variance shared with preceding layers suggested that this layer only explained *unique* variance in the model from around 220 ms. Nevertheless, it is notable that the model’s IT layer aligned with brain activity in a period that broadly coincides with the N1/N170 (or in MEG, M170) component commonly associated with orthographic processing (Bentin et al., 1999; Lin et al., 2011; Ling et al., 2019; Pleisch et al., 2019; Zweig & Pylkkänen, 2009) and which has a likely vOT source (Allison et al., 1994; Nobre et al., 1994; Pleisch et al., 2019; Woolnough et al., 2021). Overall, the hierarchical sequence of representations that emerged in CORnet-Z trained on images of words matched the progression observed in literate humans’ brains, from lower-level, more retinotopic representations of letter shapes, to more abstract, invariant representations of letter identities encoded in ordinal positions in words. However, because representations in ANNs are emergent, rather than computationally constrained by a more interpretable model, it is inherently unclear how cognitive scientists should interpret the alignment between representations that emerge in ANNs like CORnet-Z and the neural representations of humans (Guest & Martin, 2023).

We propose that optimal transport theory may provide a framework for developing a model of early orthographic, graphetic processing, which avoids unrealistic constraints on the orthographic inputs, yet is more computationally explicit and interpretable in its relationship to neural representations. Optimal transport refers to the efficient transportation of mass between two distributions, incurring a minimal transportation cost. For two points of equal mass in Euclidean space, the optimal transport plan between them can be trivially solved via their Euclidean distance. Optimal transport theory is primarily concerned with the generalisation of this problem to cases of many-to-many mapping, where we wish to efficiently transport mass between entire distributions of such points simultaneously (Peyré & Cuturi, 2020; Villani, 2009). Optimal transport solutions typically minimise *Wasserstein distance*, or "earth mover’s distance", which represents an overall cost of transporting mass, calculated as the product of masses and the distances they must travel in the transport plan. Previous approaches to calculating distances between visual shapes and letters presented to participants have often used measures of visual overlap (e.g., Dunn-Rankin et al., 1968; Fischer-Baum et al., 2017; Humphreys et al., 1988; Laws & Gale, 2002; Legros & Grant, 1916; Ling et al., 2019; Op de Beeck et al., 2008; Qu et al., 2022; Sun et al., 2018), which can capture the extent to which letters share large components (e.g., strokes). Wasserstein distance permits a more elaborated description of shape similarity, representing a more global cost for the overall shape distance (**Supplementary Materials A**) and, when applied to letters, a more complete description of the (dis-)similarity structure among letter shapes.

Accounting for global shape in this way may also better reflect the flexibility with which humans recognise the shapes of letters. For instance, literate humans routinely navigate a range of typefaces across which identical letter identities share global shape features, even though individual component features may not perfectly overlap. In the present study, we test this proposal with an RSA analysis of letter recognition processes. While previous studies have examined the representational similarities of letter and word shapes (e.g., Agrawal et al., 2019, 2020; Fischer-Baum et al., 2017; Qu et al., 2022; Rothlein & Rapp, 2014, 2017), to our knowledge, our study is the first to model these representational similarities via an optimal transport framework.

## The Present Study

In this preregistered study, we used RSA to examine the extent to which optimal transport Wasserstein distance aligns with patterns of neural activity elicited by letters, comparing its performance to that of Jaccard distance (Gilbert, 1884; Jaccard, 1901), a common measure of shape overlap, calculated as the union (e.g., area of shape overlap) divided by the intersection (e.g., total area of the overlaid shapes). Our aim was to test whether optimal transport theory provides a computationally explicit and interpretable model of the neural representation of letter shapes in humans. To this end, we exploit the high temporal resolution of electroencephalography (EEG) to constrain our hypotheses to a time window from 150-225 ms, matching the typical latency of the posterior N1 event-related potential (ERP) component associated with orthographic processing of letters and words (Bentin et al., 1999; Fraga-González et al., 2021; Lin et al., 2011; Ling et al., 2019; Madec et al., 2012; Petit et al., 2006; Pleisch et al., 2019; Rey et al., 2009; Wong et al., 2005; Zweig & Pylkkänen, 2009). We analysed patterns of neural activity elicited by letter stimuli in an alphabetic decision task (Grainger & Jacobs, 1991; Jacobs & Grainger, 1991), chosen to ensure that participants were attending to the shapes of letters.

We expected that the two measures of letter dissimilarity would share some information, and that both would align with neural representations of letters in the N1 window to some extent, but that Wasserstein distance should show greater representational alignment than Jaccard distance, as it provides a richer description of the global shape information relevant to perception of real-world letters. Specifically, we hypothesised (Hypothesis 1) that the correlation between the Normalised, Mass-Scaled Optimal Transport Wasserstein Distance matrix and neural representational dissimilarity matrices (RDMs) will be more likely to be greater than zero than it is to be less than zero, (Hypothesis 2a) that the correlation between the Normalised, Mass-Scaled Optimal Transport Wasserstein Distance matrix and neural RDMs will be higher than that between the Jaccard Distance matrix and neural RDMs, and (Hypothesis 2b) that the correlation between the Normalised, Mass-Scaled Optimal Transport Wasserstein Distance matrix and neural RDMs will uniquely explain more variance than the correlation between the Jaccard Distance matrix and neural RDMs, as assessed via partial correlations.

These hypotheses, as well as the data acquisition and analysis procedures, were preregistered^1^ on the Open Science Framework (https://osf.io/xwma2). The procedure and main analysis that we report here conform to the steps outlined in our preregistration, with the exception of two changes to the independent components analysis (ICA) used in the preprocessing pipeline. We outline those changes in the *Preprocessing* section, and explain them in more detail in **Supplementary Materials B**.

Finally, we ran exploratory analyses to gain further insight into our findings. First, we evaluate the performance of forms of Wasserstein and Jaccard distance that are invariant to geometric transformations, like translation and rescaling, reflecting the invariance known to emerge during ventral-stream visual orthographic processing. Second, we compare the performance of Wasserstein and Jaccard distance to that of two ANN models, ResNet-50 1.5 (K. He et al., 2015) and CORnet-Z (Kubilius et al., 2018), trained on the ImageNet image dataset (ImageNet Large Scale Visual Recognition Challenge 2012 dataset; Russakovsky et al., 2015) and images of letters. ResNet-50 is a 50-layer-deep convolutional ANN, widely used in computer vision tasks. CORnet-Z, which has previously been employed as a model of human word recognition (Agrawal & Dehaene, 2024, 2025; Hannagan et al., 2021), is a shallow convolutional ANN with an architecture informed by the primate ventral visual stream.

We find support for our hypothesis that Wasserstein distance aligns better than Jaccard distance with patterns of neural activity elicited by letters. We also find that implementations of Wasserstein distance that are invariant to geometric transformations better align with later brain activity, suggesting that Wasserstein distance could be useful for modeling the transition from retinotopic to more categorical perception of letter shapes. Finally, we show that while Wasserstein distance is more computationally constrained and interpretable than features derived from ANNs, the best performing variants of Wasserstein distance perform similarly to the best layers of ANNs in predicting brain activity elicited by letters. In sum, we suggest that optimal transport can provide a powerful yet interpretable framework for modeling graphetic processing of single letters.

### Methods

#### Participants

On the basis of a pilot experiment, we decided to recruit a sample of at least 15 participants. Using a very similar preprocessing and a preliminary version of the analysis pipeline to that applied here, we detected Bayes Factors of >10,000 in favour of all our hypotheses in the pilot experiment, with a sample of five participants. To examine our expected sensitivity to the hypothesised effects, we conducted 500 simulations using estimates derived from the pilot experiment, assuming our pilot participants to be representative of any future sample. Simulating datasets from 15 participants showed that all simulations had Bayes Factors >15 in the simulated direction, representing strong evidence (Jeffreys, 1961; Kass & Raftery, 1995), for all hypotheses. Moreover, the smallest Bayes Factor of any simulation was 84.1 in favour of H2a, and no simulated dataset at *N*=15 showed a Bayes Factor >15 in the opposite direction to that hypothesised for any hypothesis. Over 90% of simulations revealed Bayes Factors >1,000. We note that the five pilot participants were mostly aware of the purpose of the study. However, given the low-level nature of the effect, we did not expect this to have greatly influenced the pilot experiment. In addition to our minimum sample size of 15, we also specified a subsequent stopping rule for a scenario in which we estimated Bayes Factors which we would not have considered conclusive (i.e., <15) from our sample size of 15 participants (see preregistration for details).

The 15 participants (12 female, 3 male) ranged from 20 to 27 years of age (*M*=23.2, *SD*=2.7). All participants reported German as their first language, and reported that they had no proficiency in speaking or reading a language that uses a non-alphabetic script. Participants’ handedness was assessed via the revised short form of the Edinburgh Handedness Inventory (Veale, 2014), with participants only permitted to take part if they scored a laterality quotient of at least +40 indicating right handedness. All participants completed education to at least level 4 in the European Qualifications Framework (i.e., in Germany, Abitur). No participants reported diagnosis of any disorder that would affect their reading, and all participants reported having normal or corrected-to-normal vision. Data from one participant were excluded from analyses due to a technical error that interrupted recording (see **Supplementary Materials C** for details). An additional participant was tested instead so that we still reached the required sample size. Participants were paid at a rate of €12 per hour, and ethical approval was provided by the Goethe University Psychology and Sport Science Ethics Committee (ethics ID: 2023-43).

### Materials

Alphabetic letters comprised the 30 German letters in lower case (**Figure 1a**). These are the 26 alphabetic characters that German shares with English, plus the Umlaut and Eszett characters used in German: *ä*, *ö*, *ü*, and *ß*. Letter stimuli were presented in Arial Light font - a modified version of Arial with thinner strokes. This modified font was used to more closely match the stroke width of the false-font stimuli (explained below). This was done to prevent participants from using the width of characters’ strokes to strategically discriminate between letter and false-font stimuli without recognising letters.

**Figure 1.**
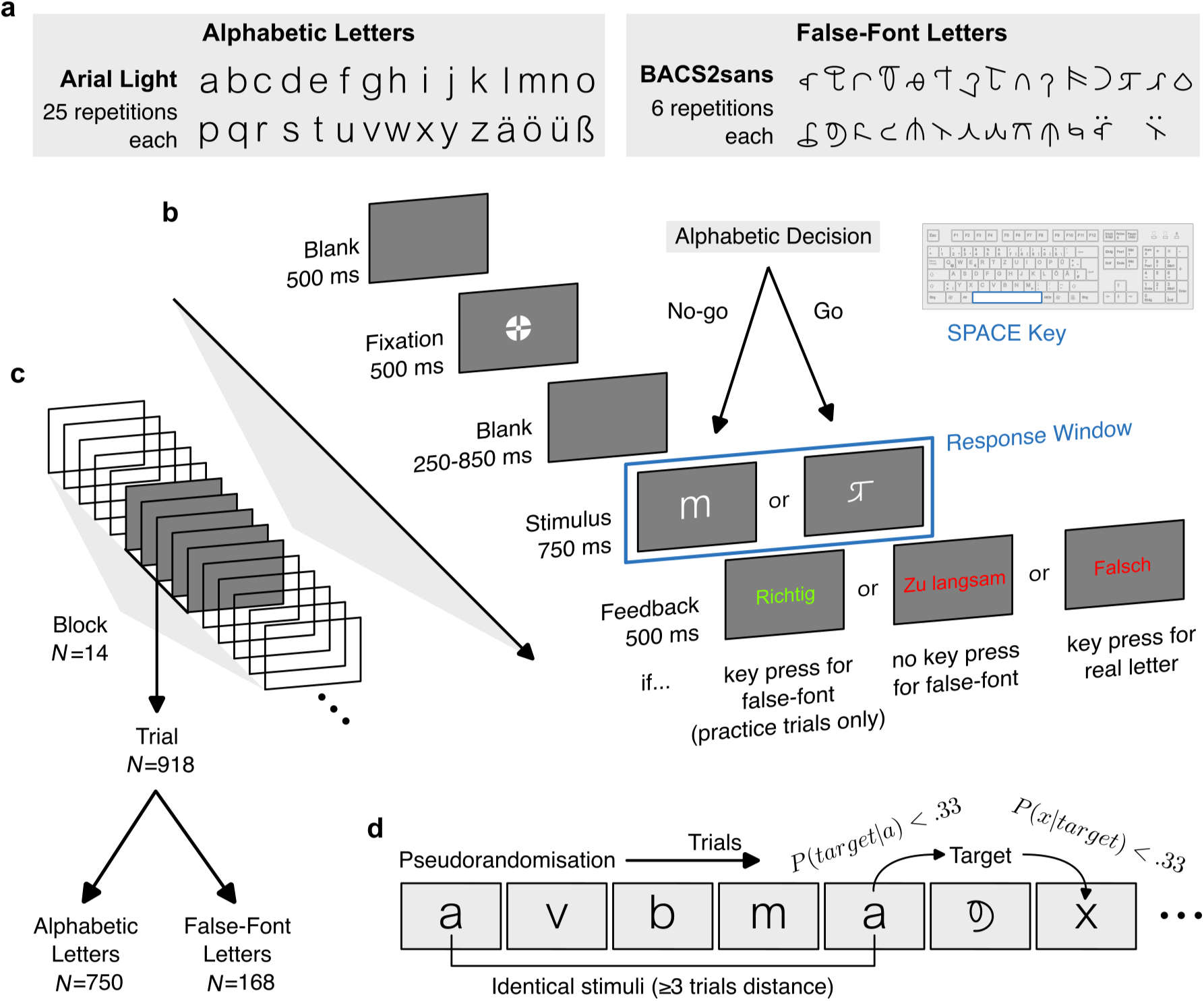
Alphabetic decision task paradigm. **(a)** The alphabetic (left) and false-font (right) letters presented to participants. **(b)** Schematic visualisation of the trial sequence. **(c)** There were 918 trials in total, across 14 blocks, split into 750 alphabetic trials and 168 false-font letters. (**d**) The order of trials was pseudorandomised to avoid stimuli being repeated and to reduce the predictabilities of target or non-target identities. Image of German keyboard adapted (recoloured) from image by A.Illigen (https://commons.wikimedia.org/wiki/File:Tastatur_DE_hellgrau.svg; CC BY 4.0).

False-font stimuli comprised lower-case characters from the BACS-2 sans-serif font (Vidal et al., 2017, **Figure 1a**). This font is designed to resemble real sans-serif letters, matched on features including the numbers of strokes, junctions, and terminations. Although we presented 30 unique target stimuli, we only used 28 matched false-font characters. The characters *ß* and *ö* are excluded because for *ß* no equivalent in BACS-2 exists, while for *ö* we observed a display error in the rendered glyph. False-font equivalents of *ä* and *ü*, formed by combining the false-font equivalents of *a* and *u* with umlaut diacritics, were displayed without error.

#### Procedure

On each trial (**Figure 1b**), participants were first presented with a blank screen for 500 ms, followed by the central "bullseye" fixation cross recommended by Thaler et al. (2013), for an additional 500 ms. There was then a blank screen for a jittered interval of between 250 and 850 ms. After the blank screen, the stimulus was presented centrally for 750 ms, during which responses to false-font stimuli would be accepted as correct. Responses were given by participants pressing the space bar on a keyboard with the index finger of their right hand, when they detected a false font character. If participants responded to an alphabetic letter, they would be shown text in red reading "Falsch" (*"Incorrect"*) for 500 ms, before the next trial began. If participants failed to respond within 750 ms to a false-font character, they would be shown text in red reading "Zu langsam" (*"Too slow"*) for 500 ms, before the next trial began.

The 30 alphabetic letters were presented 25 times each, to each participant. Each of the 28 false-font characters were presented 6 times. This resulted in a total of 918 trials, split into 14 blocks (**Figure 1c**) of 66 trials each, except for the last block which comprised only 60 trials. Each block featured 10 - 14 trials presenting false-font characters, while all remaining trials were alphabetic letters. Trial order was determined for each participant pseudorandomly (**Figure 1d**), with the constraint that no letter identity would be repeated within 3 trials of its last presentation. In this constraint, true- and false-font counterparts are considered distinct letter identities. False-font characters were also presented pseudorandomly, with the constraint that, across the entire experimental session, the maximum transition probability of any alphabetic letter immediately preceding or following a false-font character was kept <.33.

At the start of the experiment there was a practice block comprising 17 alphabetic letters and 3 false-font characters. The practice block was otherwise identical to the 14 experimental blocks. If participants failed to respond within the 750 ms time limit to >1 false-font character in the practice block, they were asked to complete the practice block again. After successfully completing the practice trials, participants were asked to repeat, in their own words, the requirements of the task, to check that they understood the experiment. During the practice trials, participants additionally received feedback for correct responses, with text in green reading "Richtig" (*"Correct"*) for 500 ms, for every false-font letter they correctly identified.

Participants were provided with self-paced breaks between blocks, during which they were presented with a summary of their performance in the block they just completed, showing: their average (median) response time in milliseconds and the number of mistakes they made together with a block-specific percentage (error rate).

Stimuli were presented at a distance of 103 cm (set via a chinrest), on a 1680*×*1050 resolution display of dimensions 47.4*×*29.6 cm, with a 60 Hz refresh rate. Letters and false-font stimuli were presented in white font on a grey-background screen, with the tallest characters’ heights subtending 1.2°of visual angle.

#### Recording

EEG data were recorded from 64 active Ag/Ag-Cl electrodes (extended 10-20 system) using a BrainProducts actiCAP system. Data were recorded at a sample rate of 1000 Hz, with an online high-pass filter of .016 Hz. Electro-oculography (EOG) data were recorded from two pairs of flat bipolar electrodes, respectively comparing voltage differences between regions to the side of the outer canthi of each eye, and between regions directly below the participant’s left eye and above the participant’s left eyebrow. Electrode impedances were kept <15 kΩ, with any electrodes that failed to reach this threshold recorded for later removal and interpolation. Data were recorded in an electromagnetically shielded booth.

Triggers were delivered via parallel port to coincide with stimulus presentation. Calibration tests with photodiodes revealed a constant delay of 8 ms between trigger delivery and change in luminance for centrally presented stimuli. Triggers were additionally used to record any key presses.

#### Preprocessing

Preprocessing was implemented using MNE Python (Gramfort et al., 2013). First, channels that had been noted during the experiment as having high impedances (*≥*15 kΩ) or being excessively noisy or flat-lining, had their voltage values interpolated (*M*=1.53 per participant; max=5 per participant). Interpolation was implemented using spherical spline models. Data were then re-referenced to the arithmetic mean of all EEG channels. We bandpass filtered data to between .1 and 40 Hz with a 4th order zero-phase Butterworth filter. Zero phase shift with effective 4th order was implemented by applying a 1st order filter in both directions, twice.

An ICA model was fit to the EEG data to identify artefacts in the data, using the extended infomax method (Lee et al., 1999) to identify 32 components. In our preregistration, we specified that we would estimate 64 components, equal to the number of EEG channels in the data. However, this led to the ICA failing to identify clear artefacts in the data, such as large blink signals, possibly due to rank deficiency. Instead, estimating 32 ICA components resulted in clearly partitioned components. As a result, we opted to estimate 32 components. We provide more detail on this in **Supplementary Materials B**.

The ICA was fit to periods of data that excluded between-block break periods, identified as periods of at least 3 seconds in duration during which no triggers were recorded, with an additional 1.1 seconds of padding after the last trigger for stimulus presentation before the break period, and .5 seconds of padding before the first trigger after the break period. The ICA was fit to a version of the data with a bandpass filter more appropriate for detecting artefactual components, using a higher high-pass cutoff of 1 Hz. In our preregistration, we specified that the low-pass cutoff for the ICA would be 40 Hz, to match the version of the data used in all subsequent analyses. However, we opted to instead use a 100 Hz low-pass cutoff, for comparability with the dataset on which the ICLabel classifier was originally trained (Pion-Tonachini et al., 2019), as recommended as best practice by the MNE-ICLabel library (A. Li et al., 2022). In **Supplementary Materials B** we explain and justify these changes in more detail. After fitting the ICA, artefactual components were identified using the ICLabel classifier (Pion-Tonachini et al., 2019), via the MNE-ICLabel Python library (A. Li et al., 2022). ICA components were identified as capturing artefacts if the ICLabel classifier assigned them any label except "brain" or "other" with a predicted probability of >85%. These artefactual components were then removed from the main version of the data filtered to within .1 - 40 Hz.

Epochs were defined as periods from stimulus onset until 1000 ms post-onset, with a 200 ms pre-stimulus baseline period. To account for the delay between trigger delivery and stimulus presentation (see *Recording*), we adjusted epochs to begin 8 ms later, to coincide with actual stimulus onset. Epochs were excluded if the participant responded to no-go (i.e., letter) trials, either during stimulus presentation, or within 250 ms of either the epoch onset or offset (including the 200 ms baseline period). Epochs were also excluded if any channels were detected as flat signals, with a minimum peak-to-peak amplitude below .5 µV, or if any channels were detected as noisy, with a maximum peak-to-peak amplitude above 150 µV. If any channels were detected as noisy or flat in more than 50% of epochs using these criteria, then preprocessing would have been re-run from the start for this participant, with the problematic electrodes interpolated along with those marked as problematic during the recording. However, this was not necessary for any participant. Finally, we verified that all participants satisfied the criteria for inclusion in the analysis concerning minimal trial numbers, as specified in the *Participants* section.

At the end of this pipeline, 1.01% of all epochs were excluded, with a mean of 7.47 epochs (range: 0-43) excluded per participant. The number of remaining epochs per letter for each participant ranged from 20 to 25 in the preprocessed dataset.

#### Main Analysis

The main analysis comprised an RSA (Kriegeskorte et al., 2008), in which RDMs were constructed for all pairs of alphabetic letters, from measures of graphemic similarity (Jaccard distance and Wasserstein distance), and from the average ERPs elicited by each letter for each participant. The correlation between these RDMs was calculated via rank (Spearman) correlation. We opted to use rank correlations, as it would allow us to capture relationships without assuming a specific link function, or that the relationships are strictly linear.

#### Model RDMs

RDMs for graphetic similarity were calculated by first drawing the 30 alphabetic letters in Arial Light font at a maximum height of 250 pixels via the Pillow fork (Clark, 2024) of the Python Image Library, and converting these to *numpy* (Harris et al., 2020) matrices. This produced matrices representing 8-bit raster images of the letters, in which the value of each element represented the intensity of a pixel value, normalised between 0 and 1 (i.e., the 8-bit intensity value divided by 255). The location of maximum overlap between the characters was then identified by means of two-dimensional cross-correlation. Specifically, the element at which peak cross-correlation was observed represented the location at which character *a* overlaps maximally with character *b* (Briechle & Hanebeck, 2001). Pairs of characters were then aligned on this basis, such that were the two matrices overlaid, the non-zero values would maximally overlap.

Jaccard and Wasserstein Distances were calculated for pairs of the resulting overlap-aligned matrices as described in **Supplementary Materials A**. Jaccard Distance was calculated as the intersection of the letters divided by their union (**Figure 2a**). Optimal Transport plans were solved via the network simplex algorithm implemented in the Python Optimal Transport library (Flamary et al., 2021), treating the pixel values as mass to be transported. Mass was scaled before solving the optimal transport plan, such that the two matrices each summed to one. The cost matrix, dictating the cost of transport between all pairs of pixels, was specified as the Euclidean distance. Wasserstein distance was calculated as the sum of the product of masses transported and the corresponding Euclidean distances (**Figure 2b**). **Figure 2c and 2d** depict the full model RDMs, the corresponding rank-transformed RDMs used in the RSA, and the relationships between them.

**Figure 2.**
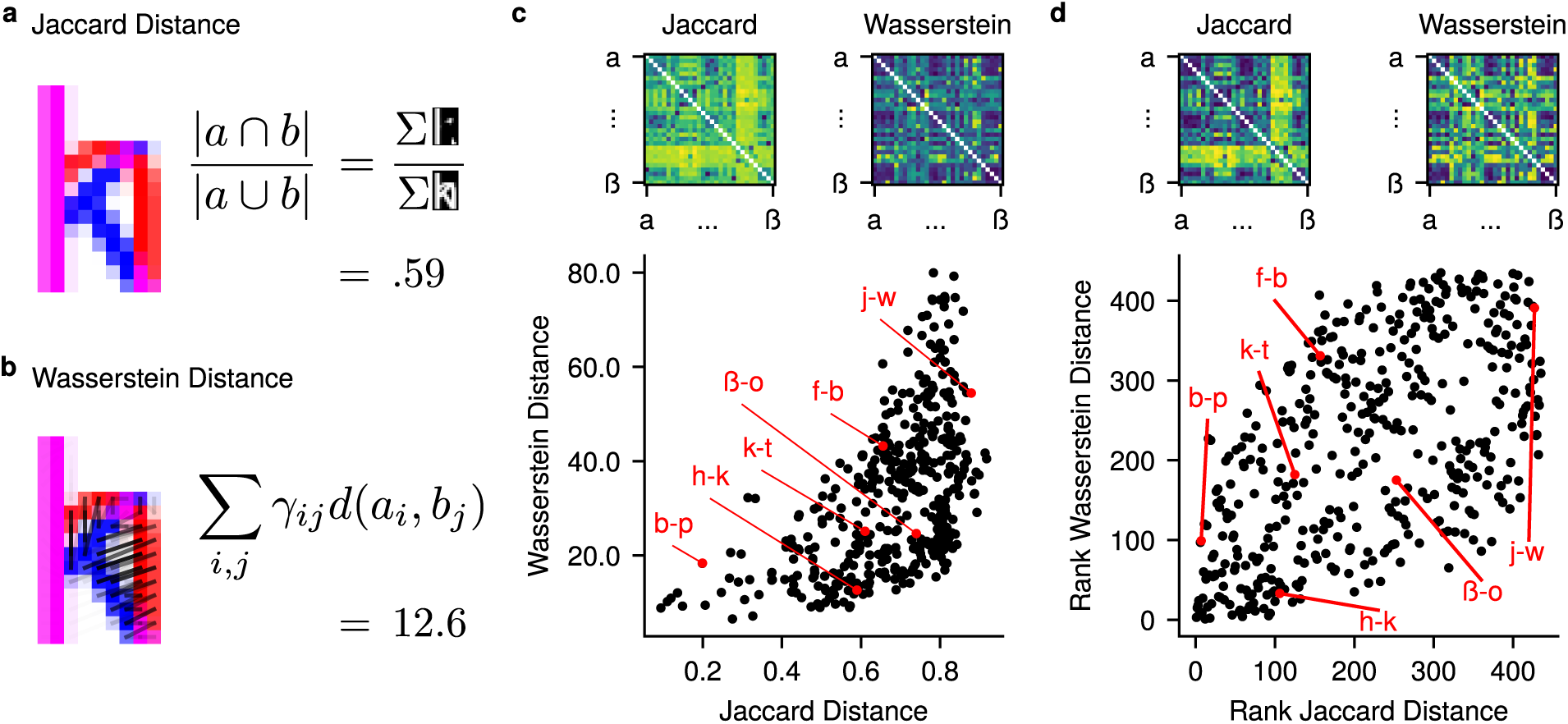
Jaccard and Wasserstein Distance RDMs. **(a, b)** Calculation of Jaccard and Wasserstein distance between two example images of Arial *h* and *k* characters (a full description is provided in **Supplementary Materials A**). For visibility, the images shown are at much lower resolution than that used in the actual analysis. **(c)** The raw Jaccard and Wasserstein distances between pairs of characters, and the relationship between their lower triangles. **(d)** The relationship between the same RDMs, but with the distances ranked from 1 to 435, reflecting the rank correlation approach used in the RSA. The rank correlation between Jaccard and Wasserstein distance is *ρ*=.57. RDMs show dissimilarities for all pairs of basic alphabetic letters in the standard order, from *a* to *z*, followed by the German characters *ä*, *ö*, *ü*, and *ß*. The diagonal is excluded from the RDMs, but would always equal zero in the raw distance values. Example pairs of letters are highlighted to show how the measures differ.

#### Neural RDMs

Neural RDMs were calculated from ERPs using the *rsatoolbox* library - a Python port and update of the MATLAB RSA toolbox (Nili et al., 2014). Specifically, a neural RDM was constructed from the Pearson correlation distances between ERPs elicited by each letter, concatenating all average amplitudes elicited by each letter end-to-end. These vectors were calculated within the time window from 150 to 225 ms (providing, at the recorded sample rate of 1000 Hz, 75 samples for each electrode). Given the posterior scalp localisation of the previously reported effects and ERPs related to orthographic processes (see Introduction section), the neural RDMs were calculated from 30 posterior electrode sites (**Figure 3a**). A neural RDM was constructed for each participant separately (**Figure 3b-d**), and rank transformed such that each value in the matrix was replaced by its rank, from 1 to 435, calculated from the lower triangle of the matrix. The noise ceiling for rank correlations with the EEG data was also calculated via the rsatoolbox library, with a leave-one-out procedure applied to participants’ RDMs (Nili et al., 2014). Specifically, each leave-one-out iteration, an estimate of the lower bound of the noise ceiling was calculated as the correlation between the RDM of the left-out participant, and average of all participants’ RDMs excluding the left-out participant. The upper bound, meanwhile, was calculated as the correlation between the left-out participant’s RDM and the average RDM across participants *including* the left-out participant. The final noise ceiling bounds were then calculated as the average of these correlations across all iterations.

**Figure 3.**
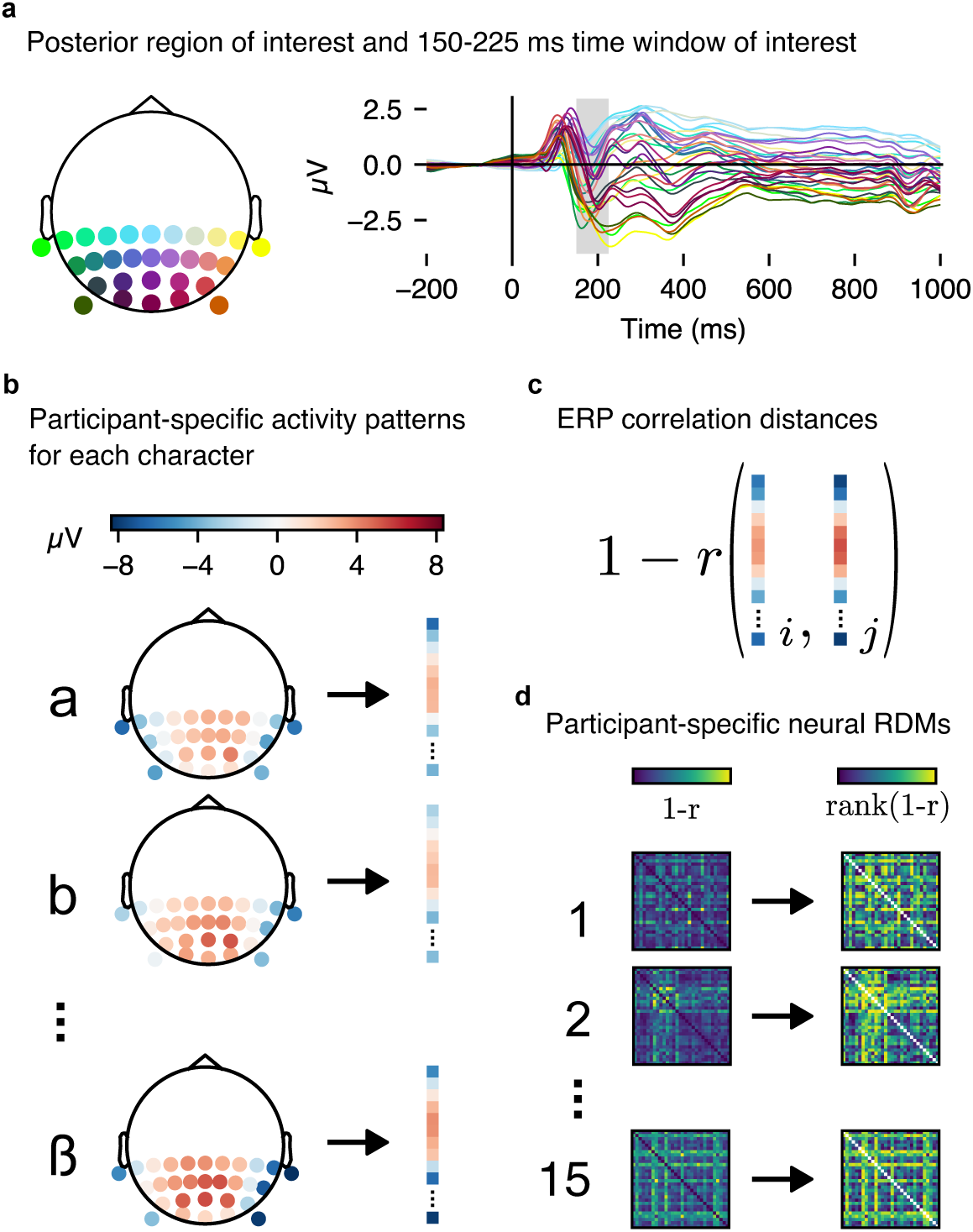
Calculation of neural RDMs. **(a)** We selected a region of interest (ROI) comprising posterior channels TP9, TP7, CP5, CP3, CP1, CPz, CP2, CP4, CP6, TP8, TP10, P7, P5, P3, P1, Pz, P2, P4, P6, P8, PO7, PO3, POz, PO4, PO8, PO9, O1, Oz, O2, and PO10. The right panel shows average ERPs of all selected channels for alphabetic letters, with the time window of interest for the planned analysis (150-225 ms) highlighted in grey. **(b)** For each participant, we calculated per-character patterns of activity as a vector of the ROI channels’ average ERP amplitudes for the given time window. Vectors from successive time points within the window were concatenated. **(c)** We calculated correlation distances between pairs of stimuli as 1 minus the Pearson’s *r* correlation coefficient of the concatenated vectors. **(d)** We calculated correlation distance matrices in this manner for all participants, and rank-transformed the values to yield the rank RDM used in the RSA.

#### Correlation Estimation

We estimated the rank (Spearman’s *ρ*) correlations between model and empirical neural RDMs (calculated from the full 150-225 ms period of interest; see previous section). The models were fit via *brms* (Bürkner, 2017), an interface to *Stan* (STAN Development Team, 2024). Full details on the models, sampling parameters, and prior specifications are provided in **Supplementary Materials E**. We additionally conducted a sensitivity analysis, which suggested that the observed pattern of effects was robust, even to priors for correlations biased very strongly towards 0, or to 1 and −1 (**Supplementary Materials F**).

In addition to rank correlations, we calculated partial rank correlations, to quantify variation in the neural RDMs that is uniquely explained by Wasserstein Distance, with Jaccard Distance partialled out, and vice versa. Posterior distributions for partial correlations were yielded by converting correlation matrices from each posterior sample to partial correlation matrices, via the *corpcor* package for R (Schäfer et al., 2021).

### Results

#### Planned Analysis

Posterior estimates (**Figure 4**) revealed a positive correlation for Wasserstein distance (median=.035, 89% HDI = [.015, .055]), and a smaller correlation for Jaccard distance (median=.014, 89% HDI = [-008, .036]). Partial correlations further showed that Wasserstein distance uniquely explained more variation in neural RDMs (median=.034, 89% HDI = [.013, .055]) than Jaccard Distance, which showed a smaller partial correlation with neural RDMs, which was likely negative (median=-.011, 89% HDI = [-.033, .013]), showing that once the variation shared by Jaccard distance and Wasserstein distance is discounted, Jaccard Distance may be systematically *anti*-correlated with neural RDMs.

**Figure 4.**
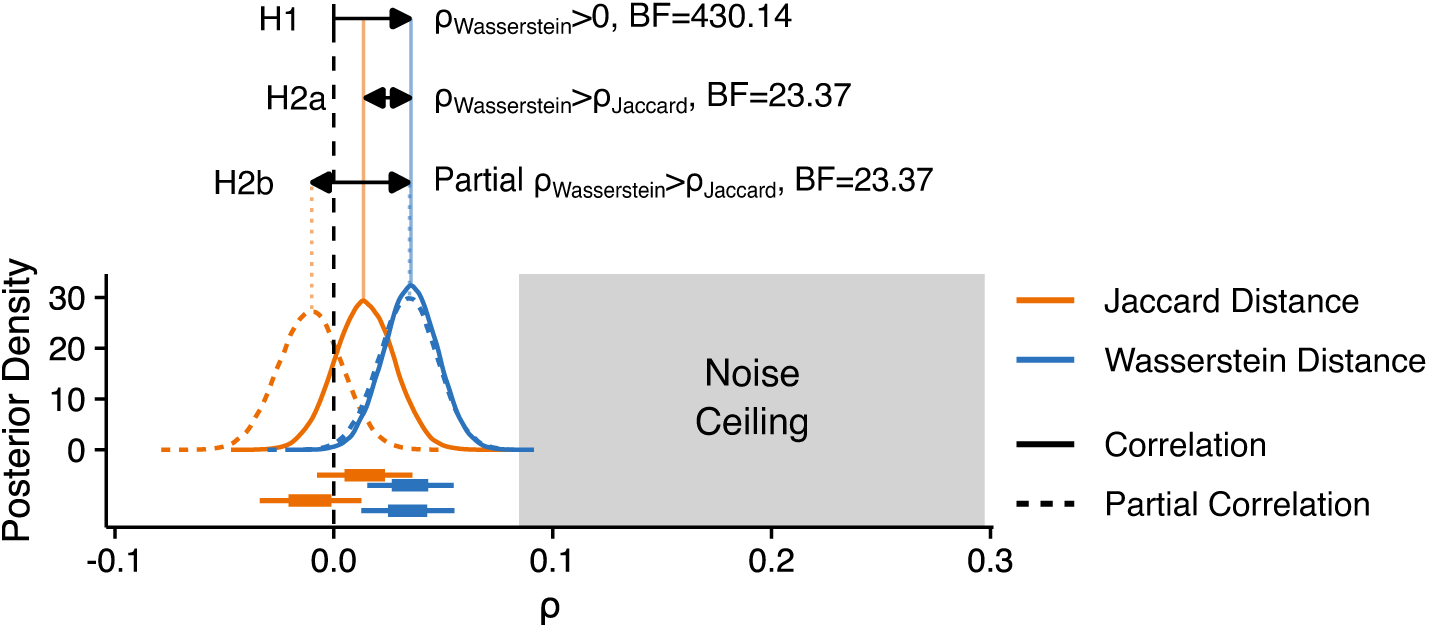
Planned analysis results. Posterior distributions for correlations with neural RDMs for posterior electrodes in the 150-225 ms window, for Jaccard (*orange*) and Wasserstein (*blue*) distance. Correlation (*solid lines*) and partial correlation estimates (*dashed lines*) are shown separately. Box and whisker plots below the densities depict the extents of the 50% (*box*) and 89% (*whiskers*) highest density intervals (HDIs) of the correlation estimates. The grey region to the right depicts the extents of the noise ceiling.

To quantify support for our hypotheses, we calculated Bayes Factors as ratios of the number of posterior samples consistent, to the number of samples inconsistent, with our hypotheses, after observing the data. This is appropriate because our symmetric priors resulted in prior odds equal to 1 for all hypotheses, such that Bayes Factors are equivalent to posterior odds (Schmalz et al., 2023). We note that the Bayes Factors we use are not tests of point hypotheses, comparing *P* (*θ* ≠ 0 | *y*) to *P* (*θ* = 0 | *y*). Instead, we test informative hypotheses (Hoijtink et al., 2008) in a directional manner, comparing *P* (*θ >* 0 | *y*) to *P* (*θ <* 0 | *y*). Were a posterior distribution for a parameter to be perfectly centred on 0, then there would be an equal number of posterior samples for both *P* (*θ >* 0 | *y*) and *P* (*θ <* 0 | *y*), resulting in a Bayes Factor of 1 exactly. A posterior distribution in the opposite direction to our hypothesis, meanwhile, would result in a Bayes Factor of *<*1.

In support of Hypothesis 1, we found strong evidence that the correlation between Wasserstein Distance and neural RDMs is greater than zero (*BF* =430.14). For H2a and H2b^2^, we also found strong evidence that the brain-model correlation was higher for Wasserstein Distance than for Jaccard Distance (*BF* =23.37).

We considered that analysing all posterior electrodes during a broad 150-225 ms window may have also captured activity related to non-orthographic processes, such as early low-level visual processing or later phonological processing. As a control analysis, we accordingly tested whether our results could have been confounded by effects of letter size, letter frequency, or phonological representations of letters. This analysis revealed a similar pattern of results for Jaccard and Wasserstein distance to that detailed above (**Supplementary Materials D**).

#### Exploratory Analyses

In addition to the planned analysis, we performed exploratory analyses delineating the time-course of brain-model alignments, implementing different forms of geometric invariance in Wasserstein and Jaccard distance, and examining how well ANNs trained on image and letter recognition align with the same neural data. We fit Bayesian models with similar specifications to those in the planned analysis (see **Supplementary Materials E** for details).

#### Time-Course Analysis

To more fully describe the dynamics of brain-model correlations, we fit time-resolved models to the full time-course of data, in bins of 10 ms. At the recorded sample rate of 1000 Hz, these correspond to non-overlapping windows of 10 samples each. As in the planned analysis, observed amplitudes from successive samples in a window were concatenated end-to-end.

Results (**Figure 5a**) showed an early peak in the correlation between brain and model RDMs at around 100 ms after letter presentation, with a similar peak correlation for Wasserstein (peak median *ρ*=.16 at 100-110 ms) and Jaccard distance (peak median *ρ*=.14 at 110-120 ms). Partial correlations (**Figure 4b**) showed, however, that Wasserstein distance uniquely explained more variation in the neural RDMs, with the partial correlation peaking at *ρ*=.11 (90-100 ms) for Wasserstein distance, but only at *ρ*=.06 (110-120 ms) for Jaccard distance. Of particular interest, the period of interest from our main analysis (150-225 ms) showed a larger correlation for Wasserstein distance than for Jaccard distance, though the difference was largest immediately before and at the start of the window.

**Figure 5.**
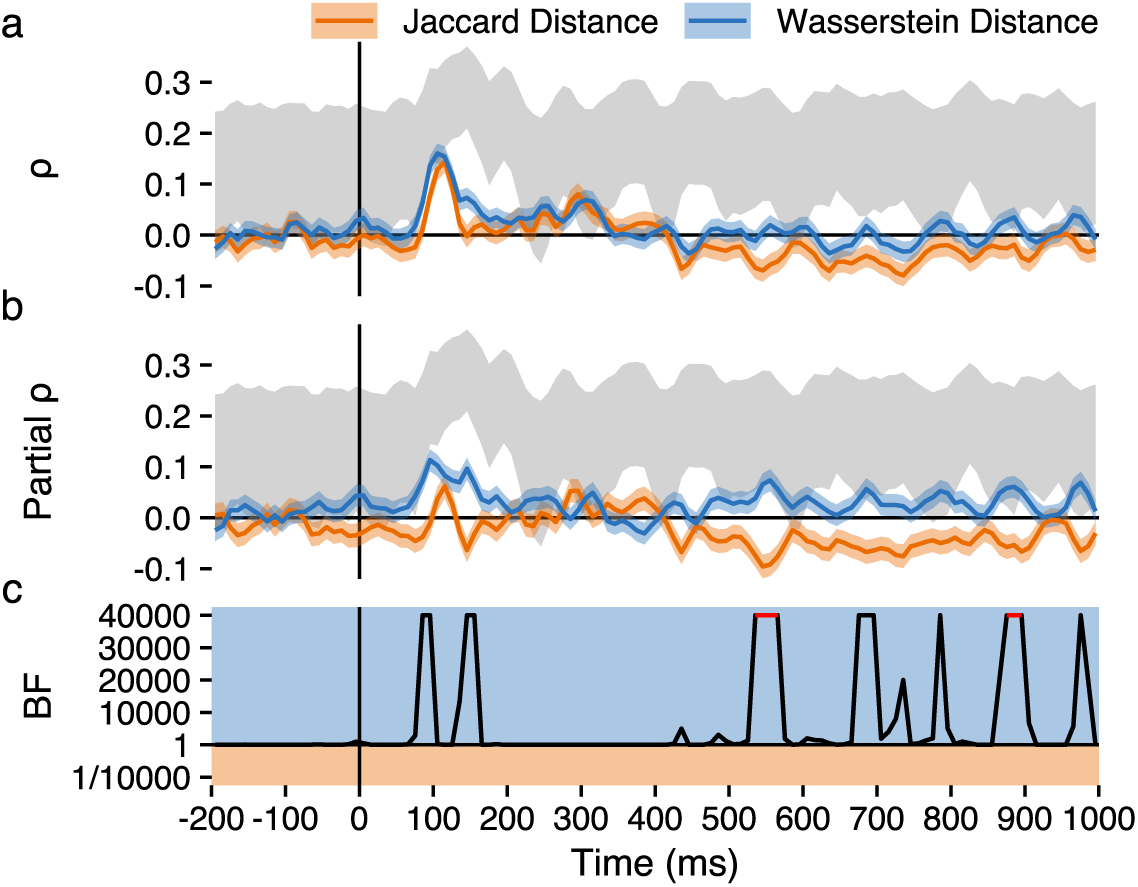
Time-resolved results. Exploratory analyses of time-resolved estimates calculated in 10 ms windows, showing point-wise medians and 89% HDIs for (**a**) rank RDM correlations, and (**b**) partial rank RDM correlations. (**c**) Time-resolved Bayes factors quantifying evidence for *ρ_W asserstein_ > ρ_Jaccard_* (Bayes factors for non-partial and partial correlations are identical). For periods in red, all posterior samples supported the hypothesis.

Surprisingly, after 400 ms, there was a sustained period during which the correlation between neural RDMs and Jaccard distance was consistently negative. This suggests that more dissimilar patterns of neural activity were elicited between letters whose shapes overlapped more.

#### Analyses of Geometric Invariance

A robust feature of the processing of visual categories across the ventral visual stream is the emergence of invariance to category-irrelevant manipulations (Grill-Spector et al., 1999; Ito et al., 1995; Rust & DiCarlo, 2010). This also applies to ventral visual processes involved in letter and word recognition, with representations becoming increasingly abstracted from visual input as information is passed along a posterior-to-anterior spatial gradient over time (Petit et al., 2006; Vinckier et al., 2007; Woolnough et al., 2021). It is across this gradient that invariance to variables like letter case (Dehaene et al., 2001; Polk & Farah, 2002) and script (Krafnick et al., 2016; Zhou et al., 2019) emerge. Similarly, letter and word processing also show a degree of invariance to *geometric* transformations - particularly to differences in retinotopic location (i.e., *translation*; Cohen et al., 2000, 2002; Dehaene et al., 2004; Dufau et al., 2008; Hannagan & Grainger, 2013; Taylor et al., 2019) and visual angle (i.e., *scale*; Chauncey et al., 2008; Eddy & Holcomb, 2009; Legge & Bigelow, 2011; Rajalingham et al., 2020). An additional geometric transformation of interest in letter perception is *rotation*, to which orthographic processes show a degree of tolerance (Fernández-López et al., 2022; Fernández-López et al., 2023; Kim & Straková, 2012; Perea et al., 2018; Yoshihara et al., 2024).

To characterise the relevance of geometric invariance to letter representations, we calculated forms of Wasserstein and Jaccard distance that are invariant to different combinations of translation, scale, and rotation. For this, we used the Nelder-Mead method of non-linear optimisation (Nelder & Mead, 1965) to identify the geometric parameters that produce the minimal Wasserstein (Figure **6a**) and Jaccard (Figure **6b**) distances between pairs of letters, varying one letter’s parameters while keeping the other constant. To avoid identifying local minima, we ran the algorithm with five starting values for each optimised dimension, spaced equidistantly from the lower to the upper bounds of the dimension. Translation was bounded between *−x* and 2*x*, where *x* is the width or height of the letter image, and zero is at the top-left origin of the image. To reduce computational intensiveness, the optimisation of translation was implemented for Jaccard distance by finding the maximum of the padded images’ two-dimensional linear cross-correlation matrices, which provides a much more efficient method for maximising overlap (Briechle & Hanebeck, 2001). Scale was optimised on a log scale, bounded between *log*(0.5) and *log*(2) (i.e., between half and twice the original size, respectively). Finally, rotation was optimised without any explicit bounds, but with starting values spaced between −180 and 180. For feasibility, letter images were drawn at a maximum height of 75 pixels, rather than the 250 pixels used in the planned analysis.

**Figure 6.**
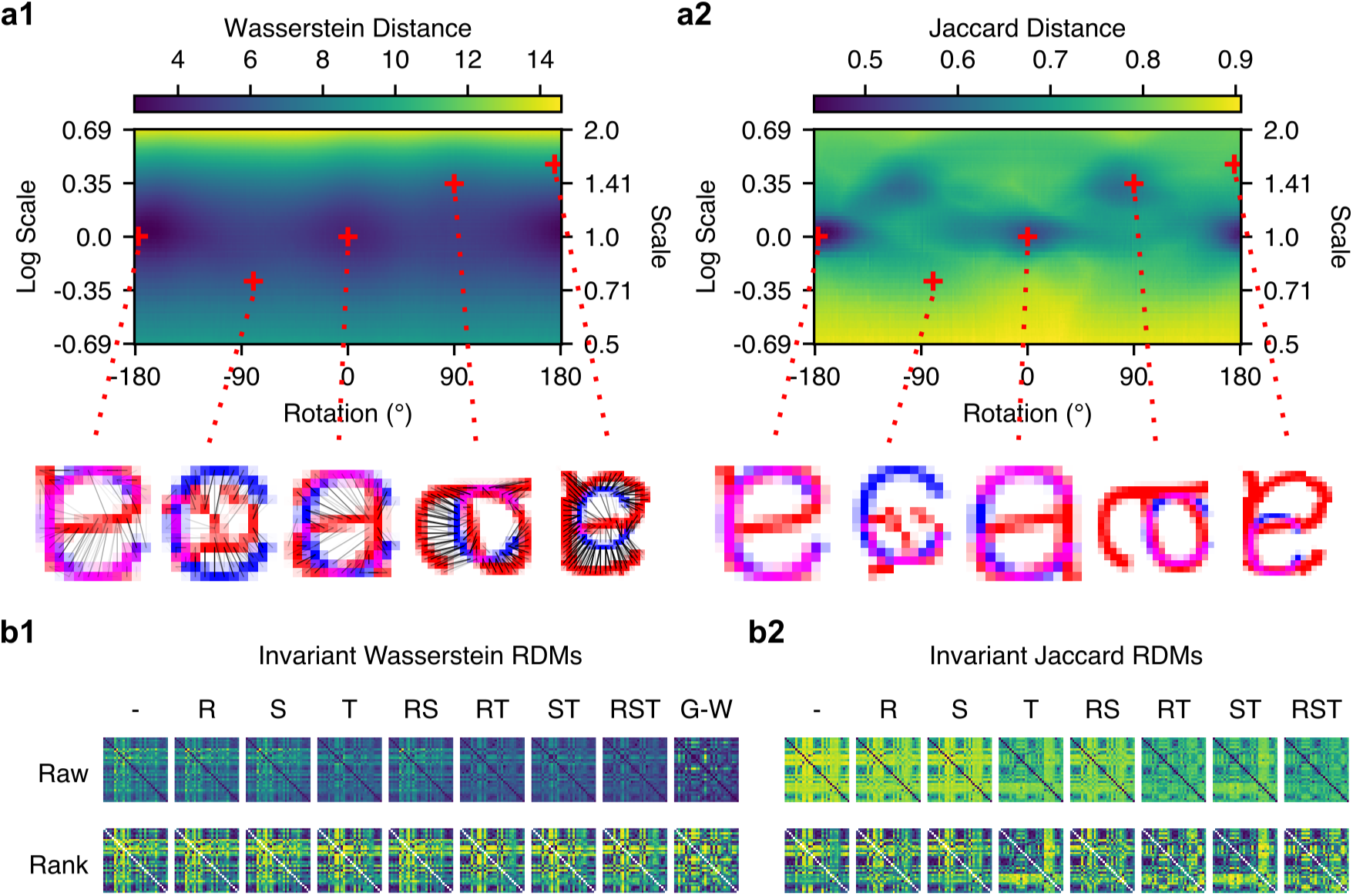
RDMs calculated with nonlinear optimisation of geometric parameters. Wasserstein and Jaccard distance RDMs were calculated with invariance to different combinations of geometric transformations. **(a1)** An example space, over which Wasserstein distance was optimised for the pair of letters, *a* and *c*. (**a2**) Jaccard distances in the same space. Translation (vertical and horizontal) was additionally optimised, via nonlinear optimisation for Wasserstein distance, and via peak two-dimensional cross correlation for Jaccard distance. Five locations are highlighted, in lower resolution than used in calculating the actual RDMs, to illustrate the differences between the spaces for Wasserstein and Jaccard distance. The example locations are identical in scale and rotation, but show differences in optimal translation. (**b1, b2**) RDMs for Wasserstein and Jaccard distance that are invariant to different combinations of geometric transformations, where values are calculated as the minima of per-letter spaces depicted in **a1** and **a2**. (*-*=default positions, *R*=rotation, *S*=scale, *T* =translation, *RS*=rotation and scale, etc.) The Gromov-Wasserstein RDM (G-W) is also depicted for comparison, though it was not calculated using the nonlinear optimisation method.

We note that in our main, preregistered analysis, we aligned characters using two-dimensional cross-correlation before calculating model RDMs. This means that the preregistered Jaccard distance measure was already translation-invariant using its optimal location, matching matrix *T*, in **Figure 6b2**, whereas the Wasserstein distance measure in the preregistered analysis was approximately translation-optimal.

As an additional comparison, we calculated Gromov-Wasserstein distance between pairs of letter images (**Figure 6b1**). This measure generalises Wasserstein distances to the problem of transporting masses between distributions that do not exist in the same metric space (Mémoli, 2011). Applied to images, Gromov-Wasserstein distance can be understood as quantifying the distances between *configurations* of mass in each image (Peyré & Cuturi, 2020). This makes Gromov-Wasserstein distance invariant to affine transformations including translation, scale, and rotation, but also less sensitive to nonlinear or non-affine transformations which preserve the configuration of mass. We calculated Gromov-Wasserstein distance via the Python Optimal Transport library (Flamary et al., 2021).

Finally, we were interested in the early peak in brain-model correlation observed at around 100 ms. As an exploratory comparison to the preregistered 150-225 ms period of interest, we examined a second, earlier period of 80-130 ms, chosen to capture this peak and reflect a typical P100 latency.

Results showed that during the 150-225 ms window, the nonlinear optimisation method of implementing geometric invariance for Wasserstein distance (**Figure 7a**), and Jaccard distance (**Figure 7b**), did not seem to greatly increase the correlation. In the 80-130 ms window, the brain-model correlation was very high for Wasserstein distance, though the most invariant measure, Gromov-Wasserstein distance (G-W), showed the lowest correlation. For Jaccard distance, there was more variability between the geometric variants, with scale-optimised Jaccard distance performing the best, though geometric invariances did not consistently improve the brain-model correlation.

**Figure 7.**
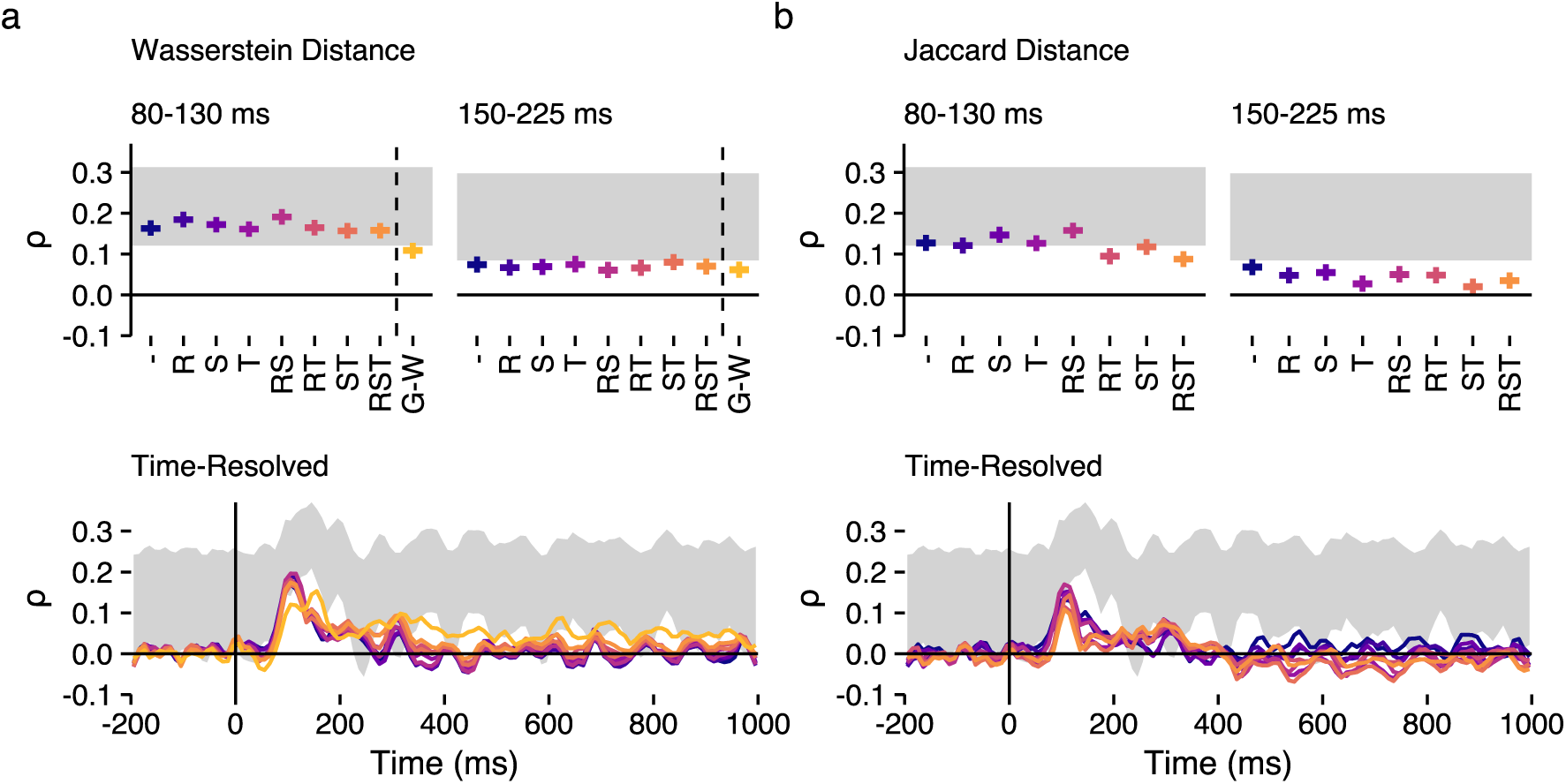
RSA results for the transformation-invariant RDMs. Spearman rank correlation estimates for transformation-invariant measures are shown for (**a**) Wasserstein and (**b**) Jaccard distance. Two windows are highlighted: an exploratory 80-130 ms window, and the original 150-225 ms period of interest. Horizontal lines depict medians of posteriors, while vertical whiskers show the extents of 89% HDIs. The lower panels show time-resolved estimates (medians of posteriors), calculated with 10 ms windows, as in the main analysis. Grey regions depict the noise ceiling, and colours are consistent across panels. As in Figure 6, model labels reflect the geometric transformations that have been optimised (*-*=default positions, *R*=rotation, *S*=scale, *T* =translation, *RS*=rotation and scale, etc.), while *G-W* refers to Gromov-Wasserstein distance.

It is notable that the RDM for Jaccard distance that was optimised for translation only (*T*, in **Figure 6b2**), which was the method used for the preregistered analysis, performed among the worst of all the Jaccard distance variants compared for the period of interest, suggesting that the variant used in the preregistered analysis may not have been representative of the other variants possible. Nevertheless, this exploratory analysis suggests that Wasserstein distance somewhat consistently outperforms Jaccard distance in predicting neural dissimilarities.

The most salient difference was observed between the transformation-optimised variants of Wasserstein distance, and Gromov-Wasserstein distance. Specifically, Gromov-Wasserstein performed somewhat worse than the Wasserstein distance measures in predicting early activity at around 100 ms (see **Figure 7a** upper panel), but performed much better in predicting neural dissimilarities in later periods, with a marked peak at around 160 ms, and prolonged period of better performance throughout the epoch (see **Figure 7a** lower panel). This is an interesting finding, and is consistent with the more invariant correspondences between letter shapes, captured by Gromov-Wasserstein distance, being processed later.

#### Comparison to ANNs

A common application of RSA is to examine the representational alignment between humans and ANNs (e.g., Caucheteux & King, 2022; Cichy et al., 2016; T. He et al., 2022; Xu & Vaziri-Pashkam, 2021). In contrast to computationally constrained measures like Wasserstein and Jaccard distance, estimates of representational similarity extracted from ANNs are largely unconstrained, instead emerging from higher-level computational goals like image classification. To compare the performance of Wasserstein distance to that of ANNs, we trained two ANN architectures on image and letter classification: ResNet-50 1.5 (K. He et al., 2015) and CORnet-Z (Kubilius et al., 2018). For both ANN architectures, we tested three variants, trained to classify images of letters in Arial Light font, images from ImageNet (Russakovsky et al., 2015), or on both letters and ImageNet images concurrently. Full details of ANN training are provided in **Supplementary Materials G**.

RDMs were calculated for ANN features by providing images of the letter stimuli as input, then extracting features via hooks attached to modules in the ANNs as implemented in the *torchextractor* library (https://github.com/antoinebrl/torchextractor). Brain-model rank correlations were estimated via multivariate Bayesian models, as described in other sections.

Results (**Figure 8**) showed that model RDMs calculated from activation patterns of the best-performing layers of ResNet-50 and CORnet-Z performed similarly to Wasserstein distance in predicting neural RDMs (compare **Figure 7**). The analysis also revealed that ResNet-50 and CORnet-Z activations aligned similarly well with patterns of neural activity, though the brain-model alignment observed for layers CORnet-Z varied much less by differences in training sets than did the alignment observed for ResNet-50. Finally, both ResNet-50 and CORnet-Z showed a pattern in which the layers which aligned best with neural activity were generally deep in the model, but not final.

**Figure 8.**
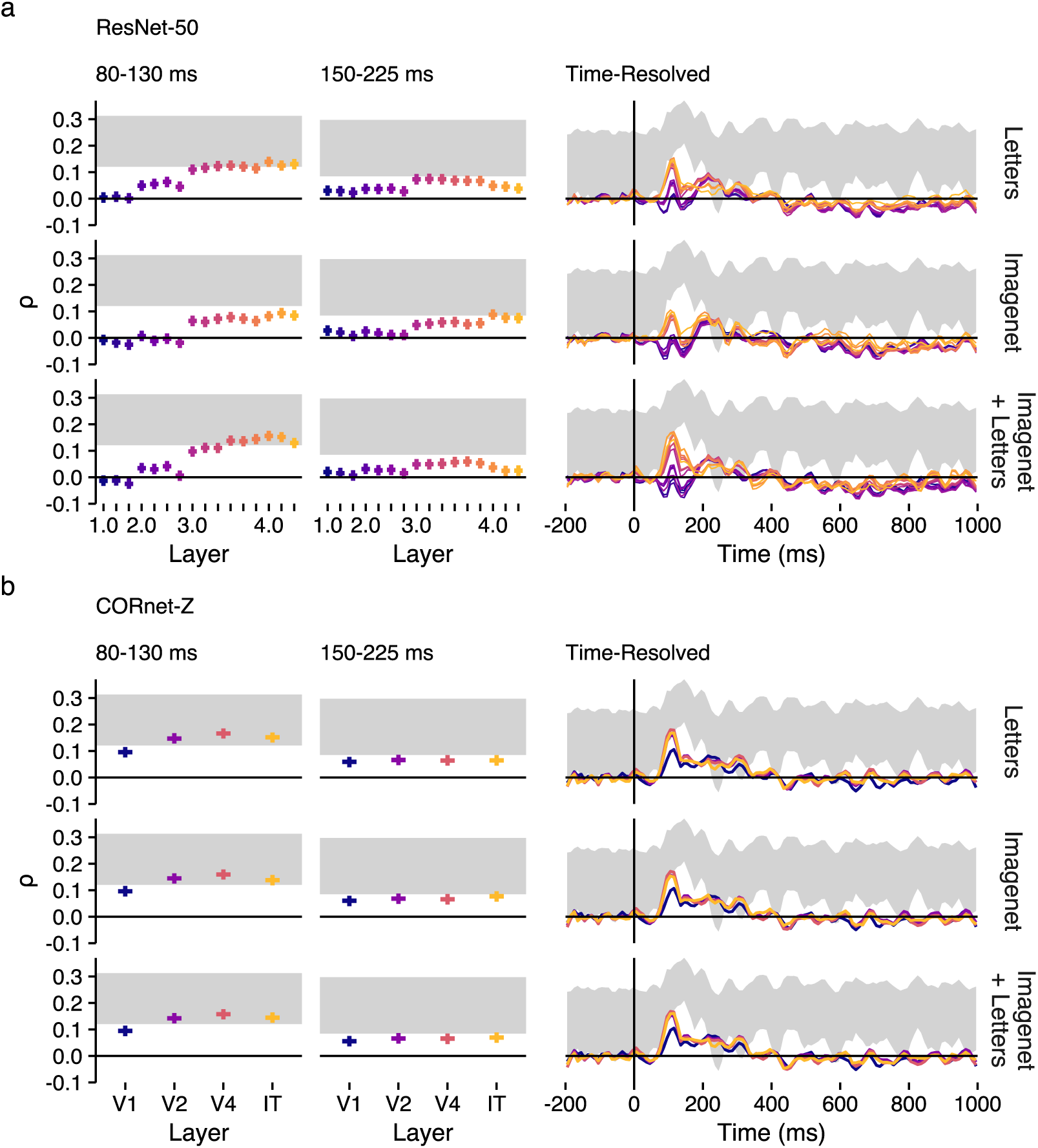
RSA results for the ANNs. Spearman rank correlation estimates for module feature similarities of (**a**) ResNet-50 and (**b**) CORnet-Z, trained on (top to bottom) letters, the Imagenet dataset, and both. The results for the two periods and time-course are calculated and shown as in Figure 7.

### Discussion

In this study we investigated whether Wasserstein distance, a metric from optimal transport theory, aligns with similarities in neural activity elicited by alphabetic letters in early, graphetic stages of processing. We demonstrated that the measure outperforms Jaccard distance, a description of simple overlap between shapes like those applied in previous models of graphetic similarities (e.g., Dunn-Rankin et al., 1968; Fischer-Baum et al., 2017; Ling et al., 2019; Qu et al., 2022; Sun et al., 2018). We additionally suggest that calculating forms of Wasserstein (and Jaccard) distance that are invariant to different geometric transformations could be valuable as a tool for investigating geometric invariances that are known to be critical in human letter and word perception. Finally, we show that Wasserstein distance performs comparably, in predicting patterns of neural activity, to less computationally explicit representations that emerge in deep artificial neural network architectures.

Our preregistered time window (150-225 ms) aimed to capture the typical latency of the N1, and our analysis revealed evidence in favour of our hypotheses during this period. However, exploratory time-resolved analyses revealed an earlier onset (around 100 ms) of the correlation between neural RDMs and both Wasserstein and Jaccard distances, and suggested that the clearest evidence for a difference between these models was in fact during earlier periods, between around 70-100 and 130-160 ms, that only partially overlapped with the earlier portion of our time window. This suggests that the low-level orthographic (i.e, graphetic) processes indexed by Wasserstein distance precede the N1, or else relate to only the earliest periods of its onset. This is consistent with existing evidence for such an early locus of more low-level, visual orthographic processing (e.g., Fernández-López et al., 2022; Ling et al., 2019; Nara et al., 2023), preceding the processing of more abstracted representations, such as letter identities and N-grams (Agrawal & Dehaene, 2025; Woolnough et al., 2021).

Investigations into visual word recognition have typically overlooked graphetic processes (Finkbeiner & Coltheart, 2009). We believe that a major reason for this has been the lack of a computationally explicit framework capable of capturing the graphetic features of letters in naturalistic typographies. Instead, graphetic similarity has often been quantified via effects on objective behaviour, such as on letter confusions (e.g., Agrawal et al., 2019, 2020; Cattell, 1886a; Mueller & Weidemann, 2012; Podgorny & Garner, 1979), or on subjective judgements, such as similarity ratings (e.g., Kuennapas & Janson, 1969; Rothlein & Rapp, 2014, 2017; Simpson et al., 2013). We argue that optimal transport theory can provide a more computationally explicit framework for describing graphetic features and their similarities. Importantly, however, we do not claim that early orthographic processing in the brain involves algorithms that solve optimal transport problems. Rather, optimal transport between images of letters *aligns* with distances between neural representations that capture the same information. At present, we remain agnostic as to the algorithms that may describe how the brain builds and processes those representations. Just as measures like Coltheart’s *N* and OLD20 have been used to establish empirical orthographic findings that computational models of visual word recognition should be able to recreate (Andrews, 1997; Norris, 2013), measures like Wasserstein distance may be used to establish graphetic-level findings that computational models of graphetic processes should be able to explain.

An advantage of applying a framework like optimal transport theory to establish such effects is that it is computationally transparent and explicit. This contrasts with the more opaque and emergent representations that can be extracted from ANNs, which we showed to perform similarly to (but not better than) Wasserstein distance in predicting patterns of neural activity elicited by letters. To be clear, we believe that ANNs can be informative tools in cognitive science, including the cognitive neuroscience of visual word recognition. ANNs provide an avenue, via alterations to model architecture and training, to formulate and test hypotheses about the nature of representations that may emerge for naturalistic stimuli. In particular, methods like RSA allow researchers to examine the alignment (Sucholutsky et al., 2023) between the representational content of ANNs, and representations reflected in behaviour (e.g., Daube et al., 2021; Marjieh et al., 2022; Rajalingham et al., 2018) or neural activity (e.g., Antonello & Huth, 2024; Caucheteux & King, 2022; Cichy et al., 2016). Researchers can attempt to infer from such representational alignment that the model and neural processes also align in computation. However, an enduring obstacle for all computational models of cognitive processes is that of multiple realisability (Putnam, 1967), which is to say that two systems that align in representational content need not align in how that representation is realised - i.e., at the level of the neural implementation or cognitive mechanisms. As Guest and Martin (2023) argue, this means that it is a logical fallacy to infer from alignment alone that ANNs implement the same computations or algorithms as the brain. This concern is not restricted to ANNs, and can be applied to inferences about human cognition made on the basis of any model, including an optimal transport description of letter similarities. Nevertheless, one route to making more valid inferences may be to constrain models to ensure that, "…models resemble phenomena because they indeed somehow capture the essence of the phenomenon" (Guest & Martin, 2023, p. 219).

Certainly, imposing theory- and biology-informed constraints on the architecture of ANNs can improve the quality of inference and understanding of cognitive and neural mechanisms that such models permit (Kubilius et al., 2018; Mano et al., 2021; Pang et al., 2021; Pogoncheff et al., 2023; Schyns et al., 2022; Sucholutsky et al., 2023). However, to borrow from Marr’s (1982) levels of analysis, ANNs constrained at the level of implementation can nevertheless remain unconstrained at the levels of computation and algorithm. Such ANNs are trained with a high-level computational goal (e.g., classification) yet researchers commonly extract and interpret features from intermediate layers, where relevant representations are emergent, rather than explicitly constrained. While model depth in ANNs trained on visual object classification does somewhat correspond to depth in the ventral visual stream (Cichy et al., 2016), the lower and intermediate model layers typically explain the most variance in neural activity, while later layers show much poorer brain-model alignment (Xu & Vaziri-Pashkam, 2021). This is a finding we replicated to some extent, with features extracted from deep, but not final, layers aligning best with neural data. A parallel can be drawn with large language model ANNs trained on next-word prediction: such models show the best alignment with neural representations of language at layers of an intermediate model depth of 60-80%, while later layers that process information more relevant to the model’s overarching computational goal show much poorer brain-model alignment (Antonello & Huth, 2024). Rather than constraining implementation or algorithm, the optimal transport description of letter similarities which we have suggested is constrained at the level of computation; it could be instantiated by multiple algorithms or implementations.

Indeed, we believe that a range of model architectures could reproduce the effect of graphetic similarity that we identified. For instance, an ANN-like model, in which letter strokes and shapes are processed by hierarchically layered feature detectors (e.g., Dehaene et al., 2005; Grainger, 2008), which show a tuning-curve response determined by their input’s distance from preferred parameters, could be expected to reproduce our finding, as Wasserstein distance between letters would likely align with distances from such tuning-curve preferences for letter detectors. Conversely, a more symbolic language-of-thought model of letter shape processing (e.g., like the proof-of-concept model implemented by Sablé-Meyer et al., 2022), in which letter shapes are encoded as minimum-length expressions in an abstracted language of geometry, could also be expected to align with Wasserstein distance between letters, as letters with more similar geometry should also be expected to possess similar specifications in such a language. Our contribution is to propose that optimal transport could serve as a model-agnostic computational framework, that can describe general principles of graphetic processing that such models share.

Such models raise the question, however, of whether graphetic distances should be calculated between pixel-based representations, as applied in our analysis, or from stroke-based representations. Both connectionist feature detectors and language-of-thought primitives may be expected to encode features of letter shapes via stroke representations, in a manner that is invariant to stroke thickness. Indeed, for arbitrary shapes, distances between topological skeletons align well with subjective judgements of similarity (Ayzenberg & Lourenco, 2019; Lowet et al., 2018) and neural correlates of shape processing in the visual cortex (Ayzenberg et al., 2022). We suggest that the optimal transport framework we have applied here could be similarly applied to stroke-based representations of letters. For instance, Wasserstein distance could be calculated from stroke-based representations, as extracted or described by existing algorithms for processing typography (e.g., Berio et al., 2022; Jakubiak et al., 2006; Knuth, 1979). We believe that comparisons between pixel- and stroke-based representations may be an interesting avenue for future work on computational descriptions of graphetic similarity. Nevertheless, a stroke-based approach would likely perform similarly well, in describing letters, to the pixel-based approach that we have applied here. This is because, unlike the shapes employed in studies on general shape processing, letters are moderately uniform in their width, such that descriptions of stroke-based representations are likely to correlate highly with descriptions of pixel-based representations.

A more general question that may be raised about the calculation of graphetic distances concerns the relevance of graphetics to visual *word* recognition. On the one hand, it has been acknowledged that whole-word shape can aid word recognition in visual scripts (Allen & Emerson, 1991; Healy & Cunningham, 1992; Lavidor et al., 2001; Perea & Rosa, 2002; Simon et al., 2007), and it is indeed possible to calculate graphetic distances for images of whole words. However, recognising words on the basis of their shape alone can be understood as an inefficient strategy, as it would require learning representations of word shapes for all words known to the reader, across variations in typography like letter case and font (Grainger, 2008). Rather than using whole-word shapes, readers are generally thought to use a more computationally efficient (Grainger, 2008, although see Pelli et al., 2003) strategy of identifying component letters via *their* shapes and features, before subsequently processing words as ordered strings of abstract letter identities (Dehaene et al., 2005; Norris, 2013). In this case, graphetic features of component letters should show downstream consequences on word recognition. Indeed, effects of visual similarities of letter substitutions have been demonstrated in orthographic priming (Marcet & Perea, 2017, 2018), and, in some cases, in unprimed lexical decision behaviour (Perea et al., 2022, 2024; Rocabado et al., 2024). A similar argument can be made about downstream effects of *typography* on word recognition. For example, words’ typographic features can facilitate or inhibit word recognition (Macaya & Perea, 2014; Minakata & Beier, 2022; Moret-Tatay & Perea, 2011), or interact with semantic processing (Kuchinke et al., 2014; Walker, 2016), and participants show online sensitivity to changes in font across successive words in lexical decision (Walker, 2008). A computationally explicit model of graphetics may provide insight into such effects of graphetics on word recognition, by quantifying, for instance, the graphetic distance associated with a specific letter substitution or font alteration.

A related question concerns the ecological validity of the letter recognition task used in this study. Explicitly requiring participants to distinguish between isolated letter and non-letter shapes may elicit processing dynamics that are not representative of naturalistic processing of alphabetic letters within words. Indeed, words are known to provide top-down context for letter processing (Cattell, 1886b; Jordan et al., 1999; Lally & Rastle, 2023; Reicher, 1969; Wheeler, 1970), and task modulates neural activity during early stages of word recognition implicated in orthographic processing, including the N1 (Chen et al., 2013; Faísca et al., 2019; Rahimi et al., 2022; Segalowitz & Zheng, 2009; Strijkers et al., 2015; Wang & Maurer, 2017). If the aim is to make inferences about reading and word recognition then it is important to consider the degree to which a given task elicits naturalistic reading and word recognition processes (Kay et al., 2023; Krakauer et al., 2017). We consider our study an important step in understanding graphetic processing of letters, but believe that future work should demonstrate a relevance to alphabetic letter processing in the contexts of word recognition and reading tasks.

Beyond alphabetic orthographies, graphetic descriptions of visual words are also relevant to modeling word recognition in more logographic scripts, in which single glyphs or characters can represent entire words. This relates more broadly to the overreliance of visual word recognition research on English and alphabetic orthographies, such that models and theories of orthographic processing show poor generalisation to other languages and scripts (X. Li et al., 2022; Share, 2008). This is despite a wealth of evidence for shared neural mechanisms of orthographic processing across scripts (Bai et al., 2011; Bolger et al., 2005; Krafnick et al., 2016; Taylor et al., 2019). A key strength of the approach to calculating graphetic similarities which we have suggested here is that it could be applied to any visual orthographic script. Given that all visual orthographies begin with a visual stage of processing, a robust framework for describing graphetic features of real-world characters could contribute to the goal of identifying universal features and mechanisms of visual word recognition processes.

Even beyond orthography, an optimal transport description may provide insights into visual shape processing more generally. Just as with letters, visual similarity is often calculated for arbitrary visual shapes via measures that capture information about overlap (e.g., Dunn-Rankin et al., 1968; Humphreys et al., 1988; Laws & Gale, 2002; Op de Beeck et al., 2008). It is possible that, as with letters, such investigations into the effects of visual shape similarity could benefit from the more nuanced and sensitive descriptions provided by optimal transport theory. Visual shape processing also shares with orthographic processing the emergence, across the ventral visual stream, of a degree of invariance in recognition to variations in geometric transformations (Goodale & Milner, 1992; Isik et al., 2014; Ito et al., 1995; Rust & DiCarlo, 2010). As a result, such research may also benefit from the invariant representations captured by descriptions like Gromov-Wasserstein distance. In sum, we view an optimal transport framework as a promising avenue for describing and investigating general visual shape processing.

A final point that we wish to mention is that our analyses have assumed that all features of a letter are equally important. However, in both letter (Fiset et al., 2008, 2009) and word (Rosa et al., 2016) recognition, the features that make up letters have been shown to be differentially informative. The optimal transport framework we have suggested may in fact be capable of accommodating such findings. First, we note that regions which best differentiate characters, and which may thus be more informative about letter identity, are likely to be those which have the greatest distance from features of other characters. For instance, the informativeness of stroke terminations (Fiset et al., 2008; Rosa et al., 2016) may be captured, in part, by the manner in which stroke terminations will be a larger distance from the image centre, where more letters are likely to have features. In this way, the present implementation of Wasserstein distance may already capture some information about the relevance of different letter regions. Second, we speculate that further differences between the perceptual informativeness of different regions of letters could be captured by distributing the letters’ masses differentially across the image space, upweighting the contribution of more informative letter regions to the overall Wasserstein distance between letters.

In sum, we provide evidence that Wasserstein distance between letters can provide a computationally explicit measure of graphetic distance between letters which aligns with the distance between corresponding neural representations. While the graphetics of real-world characters has been largely overlooked in visual word recognition research, we believe that using a framework like optimal transport to explicitly quantify aspects of letter and word shapes can provide key insights into orthographic processing, and possibly into visual shape processing more generally.

## Preregistration

The experiment, preprocessing, and planned analysis were preregistered on the Open Science Framework: https://osf.io/xwma2

## Data Availability

The EEG and corresponding behavioural data are shared openly in BIDS (Pernet et al., 2019) format via OpenNeuro: https://openneuro.org/datasets/ds005594

## Code Availability

Scripts, outputs, and a Docker container to reproduce analyses, including preprocessing, RDM calculation, and model fitting, are shared in a GIN repository: https://gin.g-node.org/JackEdTaylor/lettersim-ot-rsa

## Contributions

Contributions are described according to the Contributor Role Taxonomy (CRediT).

*JET*: Conceptualisation, Data curation, Formal analysis, Funding acquisition, Methodology, Project administration, Software, Validation, Visualisation, Writing (original draft)

*RS*: Data curation, Formal analysis, Investigation, Methodology, Software, Visualisation, Writing (review & editing)

*CI*: Formal analysis, Software, Writing (review & editing)

*CJF*: Conceptualisation, Methodology, Supervision, Writing (review & editing)

## Footnotes

1 We have made three changes to the wording of the hypotheses from our preregistration: (1) We have removed the word "partial", as partial optimal transport is equivalent to more standard *balanced* optimal transport when the masses of two distributions are equal, and we preregistered that we would scale the masses of letters to be equal, (2) we have rewordered "Wasserstein Optimal Transport Distance" to "optimal transport Wasserstein distance", to match the formatting and wording used elsewhere in this manuscript, and (3) we have reworded "neural activity" and "matrices" to "RDMs", to match standard RSA terminology. These changes do not affect how the hypotheses are interpreted or tested.

2 The Bayes Factors for for H2a and H2b are here identical, because transforming correlations to partial correlations preserves the rank order of the correlation matrix.

## Notes

This research was supported by a Fokus A|B Grant provided by Goethe University Frankfurt.

Authors report no conflict of interest.

https://osf.io/xwma2

https://gin.g-node.org/JackEdTaylor/lettersim-ot-rsa

https://openneuro.org/datasets/ds005594

### Competing Interest Statement

The authors have declared no competing interest.

### Summary of Updates

Mostly typo correction and adopting clearer wording. Also corrected statistics on the number of dropped epochs in preprocessing.

