## Supplementary Materials for "Beyond Letters: Optimal Transport as a Model for Sub-Letter Orthographic Processing"

### Contents

**A Calculation of Distance Metrics (Model RDMs) 2**

**B Changes to the Independent Components Analysis in Preprocessing 8**

**C Participant Exclusions 15**

**D RSA Analyses with Control RDMs 16**

**E Bayesian Modelling Overview 22**

**F Sensitivity Analysis 25**

**G Details on ANN Model Training and Feature Extraction 27**

### A Calculation of Distance Metrics (Model RDMs)

#### Jaccard Distance

The Jaccard index (Gilbert, 1884; Jaccard, 1901) provides a useful description of the degree of overlap between two sets. It can also be applied to describe overlap between geometric figures, where these shapes are conceived of as sets, and their composite features as set members. For instance, the measure is commonly applied in optimising and evaluating computer vision models, such as in estimating the overlap between an ideal and model-estimated bounding box. It has also been applied in neuroimaging analyses, such as in quantifying the degree of overlap between patterns of brain activation (Maitra, 2010). The Jaccard index, $J$, between two binary (e.g., black/zero vs. white/one) raster images of shapes $a$ and $b$, can be calculated simply as their intersection (the sum, or count, of the pixels in which the two shapes are overlapping) divided by their union (the sum of all pixels in shape $a$ plus the sum of all pixels in shape $b$):

$$J(a,b) = \frac{|a \cap b|}{|a \cup b|}$$

This method can also be used to describe the overlap between non-binary, anti-aliased raster images. Here, a non-binary pixel can be conceptualised as a bin in a two-dimensional histogram, capturing the proportion of pixels which, in the corresponding region of the image rendered at a higher resolution, could be expected to have a value of one rather than zero. Jaccard distance can be generalised in this way by dividing the sum of the images' element-wise minimum by the sum of their element-wise maximum. As an example, **Figure 1** shows how the Jaccard similarity between Arial letters *k* and *h* captures the overlap between their stems, while on the right of the characters, there is very little overlap between the arc of the *h* and the diagonal strokes of the *k*.

It has long been recognised that visual similarities between letters can be captured by their degree of overlap. Notably, Dunn-Rankin et al. (1968) used a variant of the Jaccard

**Figure 1**

*Jaccard Distance applied to non-binary raster images of letters.*

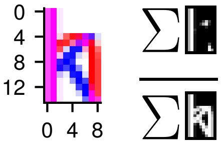


The Jaccard index is illustrated for 20-point Arial font *h* (*red*) and *k* (*blue*) characters. The letters' overlapping regions are depicted in magenta (as sums of the *red* and *blue* colour channels). The Jaccard index is calculated as the area of the letters' intersection (i.e., element-wise minimum of the matrix representations) divided by the the area of their union (i.e., element-wise maximum of the matrices), depicted in the right part of the Figure.

index[1] to calculate the maximum degree of overlap between pairs of the shapes of the 26 latin, lower-case letters, permitting translation and rotation. A factor analysis on the resulting similarity matrix reproduced categories of letters that had been found to have the highest levels of perceptual confusion for one another.

In the present study, we capture overlap via Jaccard *distance*, which is simply the inverse of the Jaccard index (i.e., $1 - J(a, b)$). To explicitly relate Jaccard distance to measures of word-level orthographic similarity like Levenstein distance, Jaccard distance can also be understood as the number of pixels that need to be inserted or deleted to change one raster image into another, normalised by the images' union (i.e., $\frac{|a \triangle b|}{|a \cup b|}$).

Related pixel-based approaches to quantifying overlap have also been used elsewhere in describing orthographic distance, such as to describe the graphetic dissimilarities of words or characters (Fischer-Baum et al., 2017; Legros & Grant, 1916; Ling et al., 2019; Qu et al., 2022; Sun et al., 2018), or to generate orthographic averages of a corpus of words (Gagl et al., 2020). However, such approaches have usually calculated similarities and distances via simple counts or sums of image pixels, unnormalised by the total area. We consider the size normalisation provided by the Jaccard Index to be a convenient

[1] In set theory terms, Dunn-Rankin et al.'s "Congruency index" can be expressed as $C(a, b) = log10\left(\frac{|a \cap b|}{|a \triangle b|}\right)$, which is equivalent to the Jaccard index transformed onto a base-10 logit scale, such that $C(a, b) = log10\left(\frac{J(a,b)}{1-J(a,b)}\right)$.

feature of the measure, as its invariance to font size reflects the invariance to size known to emerge in orthographic processing.

However, an important shortcoming of Jaccard distance is that it considers overlap in binary terms - regions of the image either overlap or do not. This neglects valuable information about the similarities between features which fail to overlap perfectly. For instance, it may be useful to capture the proximity of the left most part of the *h*'s arc to the upper diagonal stroke of the *k* in **Figure 1**. In contrast, the greater distance between the lower-right vertical stroke of the *h* and the lower diagonal stroke of the *k* may reflect the greater dissimilarity that exists between these features. Yet, according to Jaccard distance, these non-overlapping segments are equally dissimilar - as long as they do not overlap. Distances derived from optimal transport theory may provide an alternative approach that addresses this shortcoming.

### Optimal Transport Wasserstein Distance

Given two distributions of mass, $a$ and $b$, an optimal transport plan describes how to move mass from distribution $a$ to distribution $b$, incurring the minimum total cost possible under a given a loss function (**Figure 2a1**).

There exist well-developed algorithms for efficiently computing exact or approximate optimal transport plan solutions (Peyré & Cuturi, 2020). Typically, optimal transport solvers aim to minimise the Wasserstein distance (or "Earth mover's distance"), $W$, between distributions $a$ and $b$, calculated as

$$W(a,b) = \sum_{i,j} \gamma_{ij} d(a_i, b_j)$$

In our application, loss function $d(a_i, b_j)$ gives the pairwise Euclidean distances[2] between each sample $i$ in distribution $a$, and each sample $j$ in distribution $b$ (**Figure 2a2, 2b2**).

[2] $i$ and $j$ represent the *flattened* indices of the two distributions of mass, to match the dimensions of transport plan $\gamma_{ij}$, but the distance function $d$ calculates distances using samples' original 2D indices.

**Figure 2**

*Optimal transport applied to raster images of letters.*

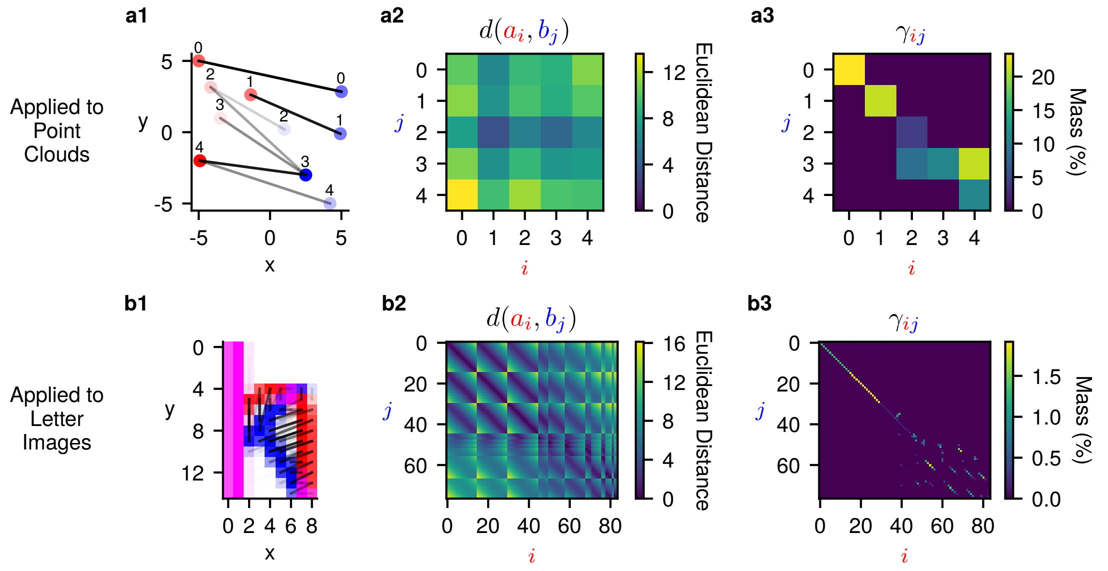


**(a1)** An example transport plan between two point clouds, where the opacity of points represents their relative masses. The opacity of the black lines between points represent the amount of mass moved from each source (*red*) to each target (*blue*) sample. Numbers above points correspond to $i$ and $j$ indices in the distance and transport plan matrices. **(a2)** The distance matrix $d(a_i, b_j)$, between each source location $i$ (x axis)and each target location $j$. (y axis). **(a3)** The optimal transport plan $\gamma_{ij}$, showing the amount of mass moved from each location $i$ to each location $j$. Panels **(b1-3)** depict equivalent information for the optimal transport between images of mass-scaled letters *h* and *k*, in 20-point Arial font.

This is multiplied by corresponding elements in a matrix $\gamma_{ij}$, representing the optimal transport plan (**Figure 2a3, 2b3**). In this matrix, values represent the amount of mass transported from each sample in $a$ to each sample in $b$. For pairs of samples where no mass is transported in the optimal transport plan, the value will be zero, such that the distance between those samples will not contribute to the Wasserstein distance. Correspondingly, pairs of samples that already perfectly overlap will be at a distance of zero from one another, similarly zeroing out the cost of transport between those samples. If it minimises the total cost, mass from one source sample $i$ can split into two target samples, $j$, or vice versa. This would be represented in the transport plan as non-zero values in two or more rows of column $i$, or columns of row $j$, respectively. Applied to describing the similarity between two raster images of shapes like letters, mass can be represented by the intensity of pixel values (**Figure 2b**; Rubner et al., 2000). Pixels can then be treated as bins in a two-dimensional histogram of mass.

A key question when using an optimal transport approach is how to deal with distributions of different mass. For example, the sum of pixel values in the letter *M* exceeds by far the sum of pixels in a smaller letter of equal font size, like *I*. One solution could be to calculate the *partial* Wasserstein distance, via a transport plan constructed for the total mass in the smaller character only. However, this would leave the mass in the remaining pixels of the larger letter unaccounted for, and would imply zero distance between the letters *I* and *M*, if the *I* overlaps with one of the vertical strokes of *M*. Other solutions include computing variants of *unbalanced* optimal transport in which an additional parameter dictates a trade-off between transportation costs and mass that the solution fails to transport, which can be conceived as a method for excluding outlying samples where the cost of transport in a given application may be greater than the cost of failing to transport the mass (Séjourné et al., 2023). However, we opted for a simple method, whereby the images of letters were mass-scaled prior to calculating Wasserstein distance. Specifically, we divide the value of each pixel by the sum of mass in the image, such that values represent the proportion of mass in each location:

$$a_i \leftarrow \frac{a_i}{\Sigma_k a_k}, b_j \leftarrow \frac{b_j}{\Sigma_l b_l}$$

As in our preregistration, proportion scaling could be described equivalently as a combination of upscaling prior to calculating Wasserstein distance, and subsequent mass normalisation of the Wasserstein metric. For instance, if the total mass of $b$ exceeds that of $a$, the mass of $a$ would be first upscaled to match that of $b$:

$$a_i \leftarrow \frac{a_i}{\Sigma_j b_j}$$

Following this, the Wasserstein distance between $a$ and $b$ could be normalised by the total mass transported:

$$W(a,b) = \frac{\sum_{i,j} \gamma_{ij} d(a_i, b_j)}{\sum_{i,j} \gamma_{ij}}$$

A final consideration may be that researchers may wish to meaningfully bound Wasserstein distance estimates between 0 and 1, such as to implement invariance to font size, like that inherent to Jaccard distance. In our study, this would have been redundant, as we rank-transform all variables, and our approach preserves the rank ordering of letters with changes in font size. Nevertheless, we suggest that one approach could be to normalise by the maximum distance in the space. For instance, where $x$ is a vector of the widths of all images, and $y$ is a vector of the heights, while $a$ and $b$ have been proportion-scaled as described above, distance-normalised Wasserstein distance could be calculated as

$$W_{norm}(a,b) = \frac{\sum_{i,j} \gamma_{ij} d(a_i, b_j)}{\sqrt{max(x) \cdot max(y)}}$$

Whereas Jaccard Distance captures how much of the letters fail to overlap, Wasserstein Distance additionally captures the distances between regions that do not overlap, accounting for more global shape information. We expected that this richer description would allow Wasserstein Distance to better predict humans' orthographic representations of letters. While optimal transport has been applied to problems of shape matching (e.g., Su et al., 2015), string matching (e.g., Tam et al., 2019), and most recently as a way of establishing mappings between representational spaces (e.g., Aoun et al., 2024; Kawakita et al., 2024; Takeda et al., 2024), to our knowledge, we are the first to apply optimal transport to describe directly the perceptual or representational distances between visually presented shapes.

### B Changes to the Independent Components Analysis in Preprocessing

We made two deviations from our preregistration in the Independent Components Analysis (ICA) step of our preprocessing: (1) we changed the number of components estimated from 64 to 32, and (2) we changed the low-pass filter applied prior to fitting the ICA. Change 1 was made because the extended infomax ICA algorithm failed to fit due to rank deficiency when we estimated 64 components from 64 EEG channels. Change 2 was made because of a discrepancy between our original low-pass filter, and that applied to the data that ICLabel was trained on. As we outline here, we believe that these changes were appropriate for the analysis.

#### 1. Change to the Number of Components

We preregistered that we would fit an extended infomax ICA with 64 components, but the ICA produced rank-deficient fits when estimating this number of components. This was accompanied by warnings from MNE for each participant, about the ratio between components' variances, for example:

```
RuntimeWarning: Using n_components=64 (resulting in n_components_=64) may lead to
    an unstable mixing matrix estimation because the ratio between the largest
   (49) and smallest (3.8e-29) variances is too large (> 1e6); consider setting
   n_components=0.999999 or an integer <= 64
```

In **Figure 3**, we show components' topographies, and a section of the time courses for estimated latent sources, for one participant (*sub-12*), from a 64-component ICA fit. This participant was chosen as they had clear blinks, with somewhat quite consistent timing after a stimulus was shown, so that blink artefacts are visible at frontal sites in the event-related potential (ERP). In **Figure 4** we show the same information when the same data are partitioned into 32 components. The ICA fit with 64 components accounts for a low proportion of variance in the EEG data overall and fails to identify clear ocular artefacts. In contrast, using 32 components greatly improves the ICA fit, including a component which captures the ocular artefacts very well and with high specificity.

Removing ICA components (identified via ICLabel with an 85% probability threshold) from the the 64-component ICA removed most of the signal from the ERPs (e.g., **Figure 5**). In contrast, identifying and removing ICA components from the 32-channel fit removed artefacts from the data while preserving non-artefactual aspects of the ERP. In sum, we considered the preregistered number of components to estimate in the ICA was inappropriate for cleaning the data, and we opted instead to estimate 32 components to avoid rank deficiency.

**2. Change to the Low-Pass Filter**

In our preregistration, we specified that the copy of the data used to fit the ICA would be first bandpass filtered to between 1 and 40 Hz. However, the ICLabel classifier (Pion-Tonachini et al., 2019), which we used to identify artefactual components, was trained to classify ICA components from datasets bandpass filtered to between 1 and 100 Hz. Requesting classifications from ICLabel when the data are filtered to between 1 and 40 Hz also produces a warning from the MNE-ICALabel (Li et al., 2022) Python library:

```
RuntimeWarning: The provided Raw instance is not filtered between 1 and 100 Hz.
    ICLabel was designed to classify features extracted from an EEG dataset
    bandpass filtered between 1 and 100 Hz (see the 'filter()' method for Raw and
    Epochs instances).
```

We therefore deviated from our preregistration, applying a 1-100 Hz bandpass filter to the copy of the data used to fit the ICA, so that our data were more similar to those that the ICLabel classifier was trained on. This change did not greatly affect the results. First, the ICA fits produced very similar results, regardless of whether a 100 (**Figure 4**) or 40 Hz (**Figure 6**) low-pass cutoff was used. Removing components identified via ICLabel also resulted in very similar ERPs regardless of which cutoff was used (**Figure 7**). Nevertheless, we decided to address the warning, changing the low-pass cutoff to 100 Hz, for consistency with ICLabel.

**Figure 3**

*First 20 ICA components for participant sub-12 when ICA is fit with 64 components, after bandpass filtering to 1-100 Hz.*

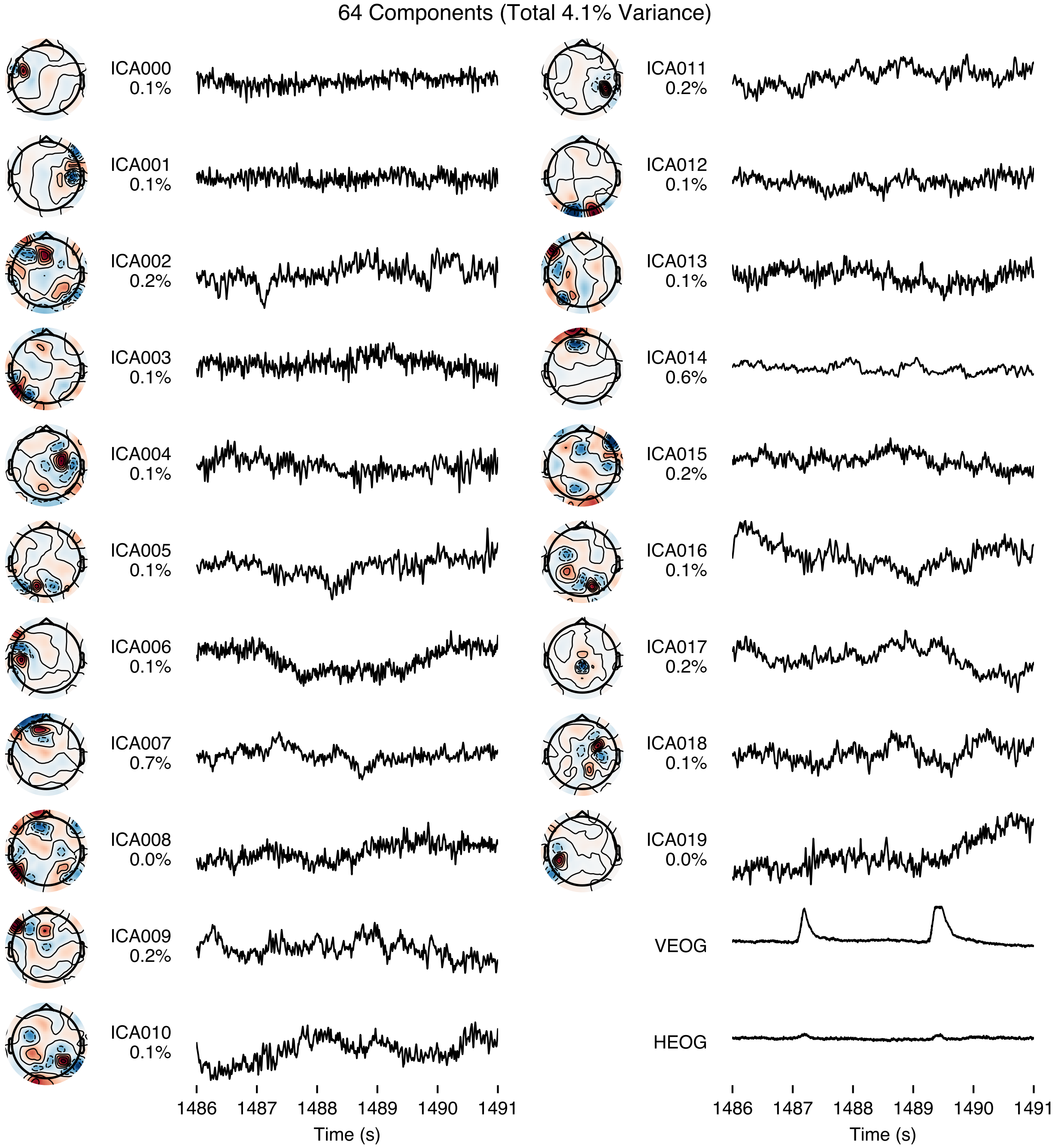


Only the first 20 components are shown. For each component, topographies are shown on the left. Time courses on the right show latent sources for an example 5 second segment during which two blinks were observed. Percentages reflect estimated proportion of variance in EEG data explained by each component. For comparison, the last two time courses depict recorded signals from the vertical and horizontal electro-oculography (EOG) electrodes.

**Figure 4**

*First 20 ICA components for participant sub-12 when ICA is fit with 32 components, after bandpass filtering to 1-100 Hz.*

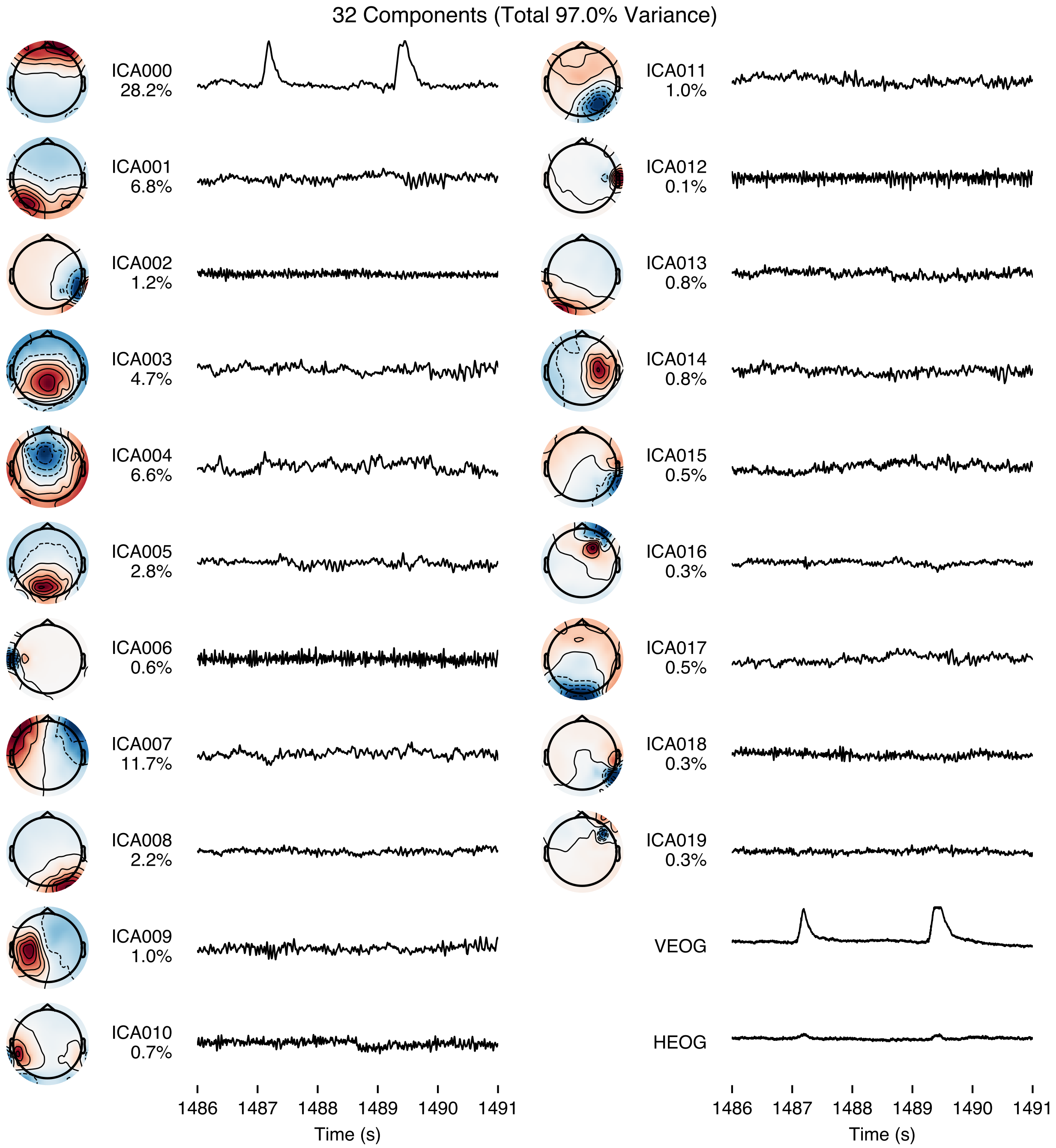


See **Figure 3** for details.

**Figure 5**

*Comparison between ERPs after removing problematic components with ICLabel from 32- and 64-component ICA fits.*

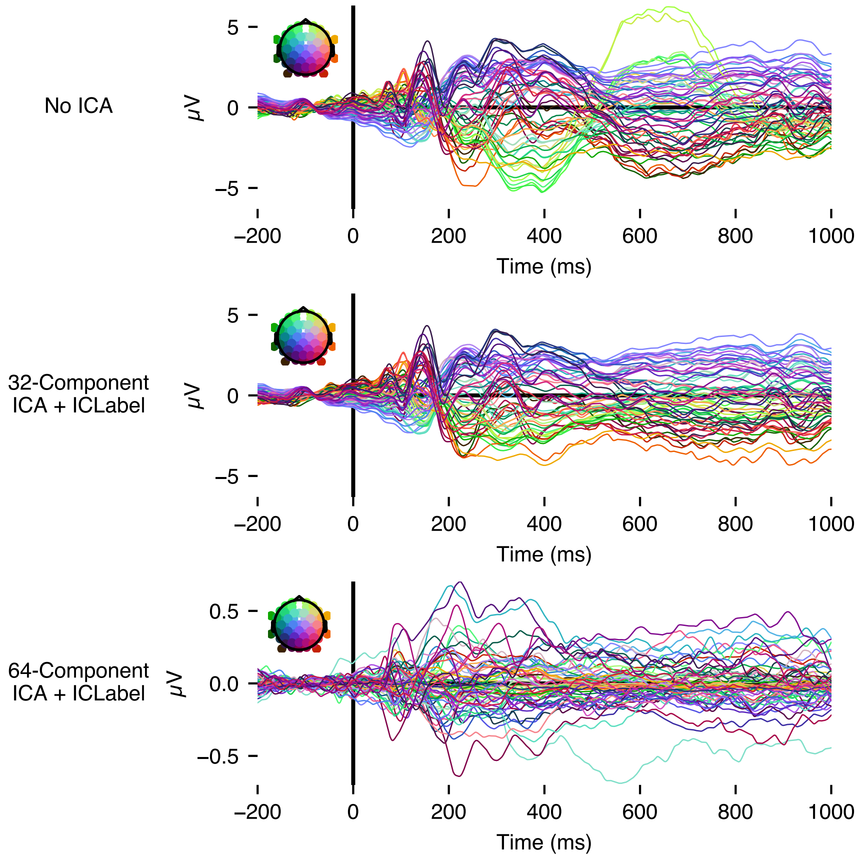


Butterfly plots show average ERPs for all alphabetic letters, from the same participant as shown in **Figures 3 and 4**. When ICLabel is applied to the 32-channel fit, most of the ERP is unchanged, except for a frontal peak at around 600 ms from average time-locked blink activity. For the 64-channel fit, meanwhile, most variance is removed from the ERP (*Note:* the y limits in the 64-component panel are reduced for visibility).

**Figure 6**

*First 20 ICA components for one participant when ICA is fit with 32 components, after bandpass filtering to 1-40 Hz.*

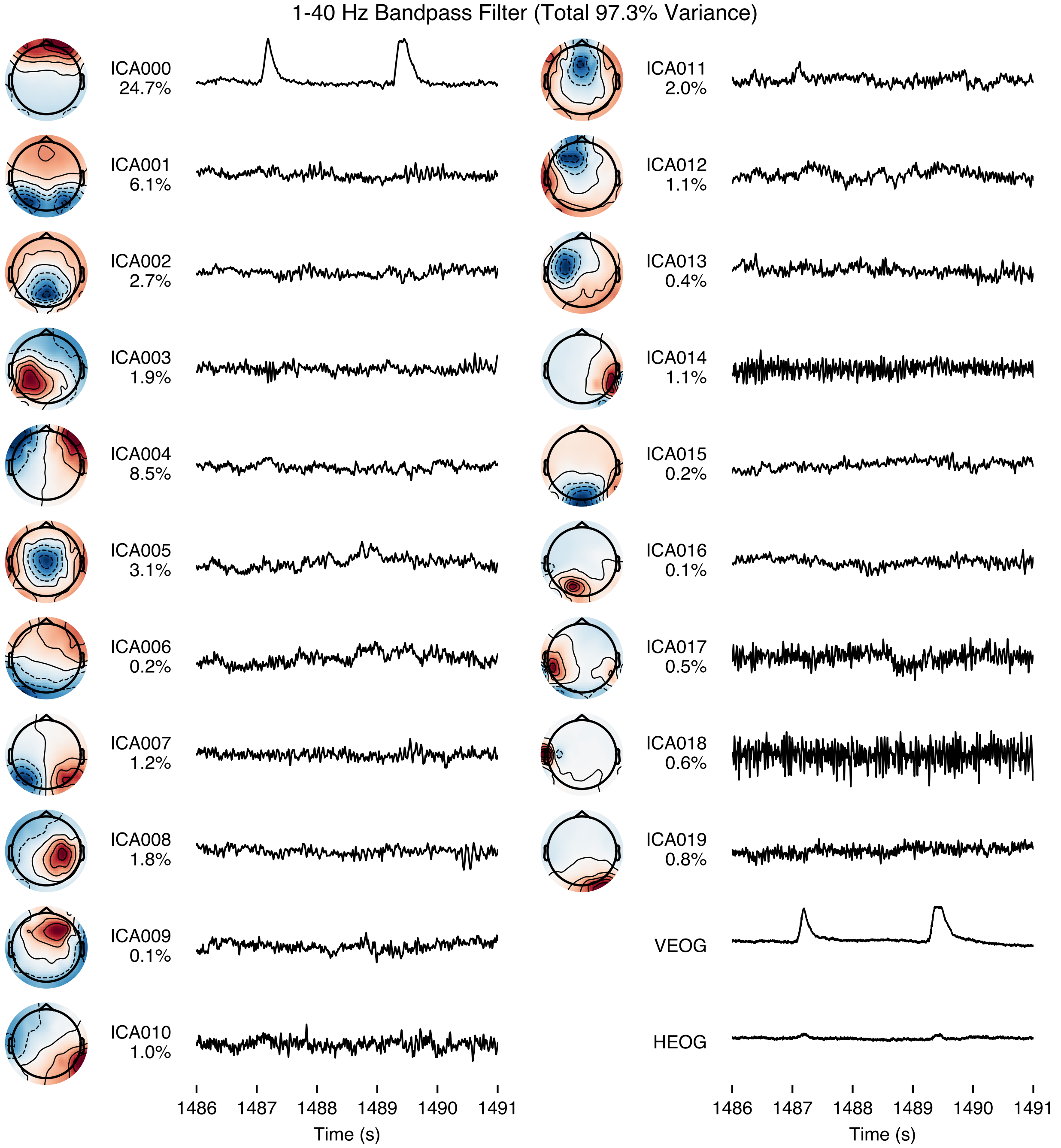


See **Figure 3** for details. The ICA fit is very similar to that shown in 4, where a 1-100 Hz bandpass filter was applied. Importantly, the percentages refer to variance explained in the copy of the data filtered for the analyses, with a .1 - 40 Hz bandpass.

**Figure 7**

*Comparison between ERPs after removing problematic components with ICLabel when the ICA was fit to data with a 1-40 versus 1-100 Hz bandpass filter.*

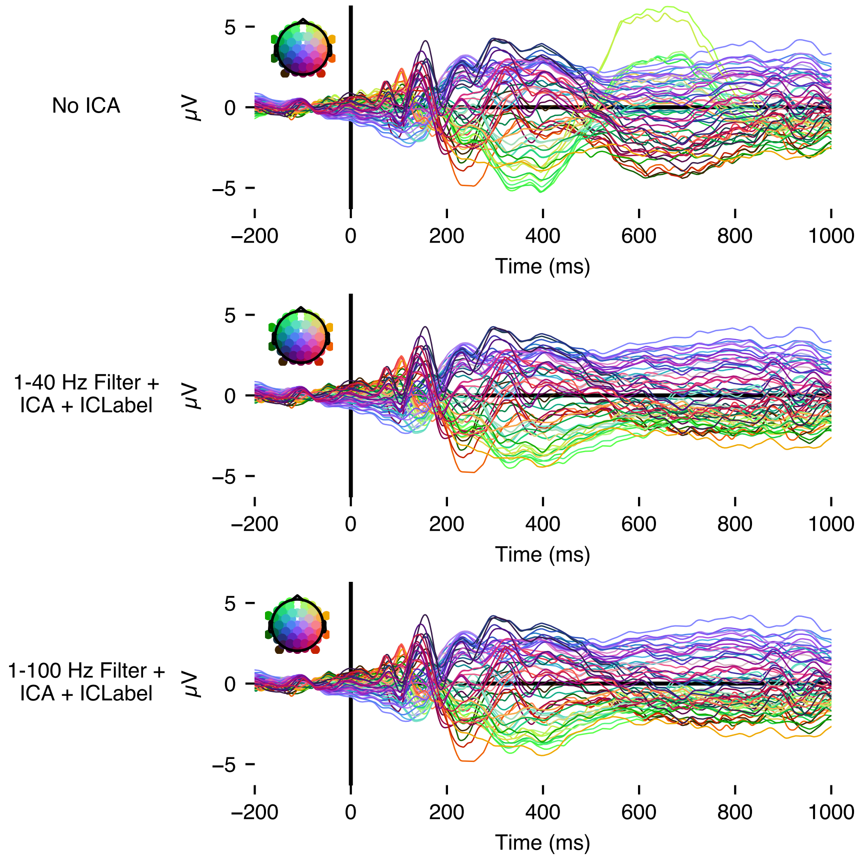


Butterfly plots show average ERPs for all alphabetic letters, from the same participant as shown in other figures in this section. Notably, the results are almost identical.

### C Participant Exclusions

We specified in our preregistration that we would exclude participants who make more than 50 errors in their behavioural responses. In our paradigm, errors could refer to a failure to respond to a target trial within <750 ms, or incorrectly providing a response to a non-target trial. We also specified that we would exclude participants who had more than 8 EEG channels marked as "bad" during the experiment, due to excessive noise or impedances >15 $\mathrm{k\Omega}$. Finally, we specified that we would exclude participants if, after the trial exclusions specified in preprocessing (see Preprocessing section of manuscript), there were fewer than 10 epochs for any of the 30 stimuli we presented. No participants were excluded according to these criteria.

However, for one participant (not counted in the 15 participants described and analysed in the manuscript) we experienced an unexpected fault in the EEG recording that prevented the recording from being saved after the first block had been completed. This meant that for this participant, the experiment had to be restarted from the beginning after the first block had already been completed once. We do not analyse this participant's data in our analyses, but the data are included in the data repository associated with the manuscript.

### D RSA Analyses with Control RDMs

We considered that confounding variables could underlie the pattern of results we observed for Wasserstein and Jaccard distance. In particular, previous findings have suggested that low-level visual and phonological information show an early locus of effects in word perception that overlaps with orthographic processing (Ling et al., 2019). We considered that letters' (1) visual size, (2) orthographic frequencies, or (3) phonological mappings may have correlated with the distance measures we calculated. This may have confounded our interpretation of the RSA results. To investigate this, we fit the same multivariate Bayesian models as specified in the main analysis, but with additional representational dissimilarity matrices (RDMs) for these features (**Figure 8**). In this way, we could calculate partial correlations to check whether the brain-model correlations for these possible confounds could account for the pattern of effects we observed.

#### 1. Visual Size

We considered that the size of the letters may have influenced the RSA results. We expected that this may be particularly relevant to the earlier components of the stimulus-evoked event-related potentials (ERPs) that we were primarily interested in. We first calculated visual size for each letter as the sum of its pixels. Visual size RDMs were then calculated as the differences between pairs of letters' pixel sums.

#### 2. Orthographic Letter Frequency

We use orthographic letter frequency to refer to the frequency with which letters occur in texts. To calculate this for German, we counted the occurrence of letters in the SUBTLEX-DE corpus (Brysbaert et al., 2011) of word frequencies. Specifically, letter frequencies were calculated weighting each occurrence of a letter in a word by the frequency of the word it occurs within. For instance, the letter *a* occurs twice in the word *ausgebrannt* ("burnt out"). The word *ausgebrannt* is observed 44 times in the SUBTLEX-DE dataset, and the letter *a* is observed twice in each occurrence, so this

**Figure 8**

*Control RDMs and the features they were constructed from.*

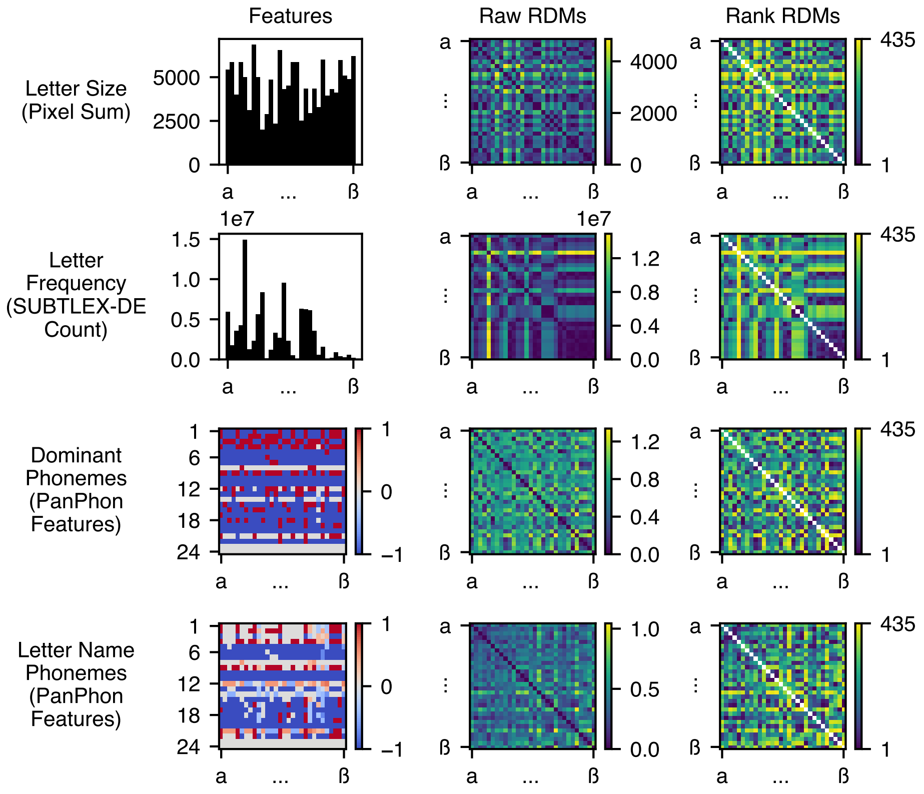


(Left) features, (centre) raw RDMS, and (right) rank RDMs for the four control variables. Latin letters are ordered according to their standard alphabetic order (*a* to *z*), followed by the German characters in the order *ä*, *ö*, *ü*, and *ß*.

word contributes a letter frequency count of 88 to that letter. We simply summed these frequency-weighted counts for all lower-case letters. To calculate the orthographic letter frequency RDMs, we then calculated the differences between the counts for all pairs of letters. Upper-case letters (e.g., at the start of every German noun) were not counted.

#### *3. Phonological Mappings*

We considered that, while reading letters in isolation, participants may activate phonological representations for the letters' (a) most common corresponding phonemes, or (b) names.

#### *(a) Dominant Phonemes*

We considered that, when presented a letter, participants may have activated representations for the phoneme that the letter usually represents in German. For instance, in German, the letter *z* usually represents the voiceless alveolar sibilant affricate, [t͡s]. On the one hand, letters' pronunciations are altered by the context of surrounding letters, reflecting phonotactic patterns. For example, the letter *c* is usually pronounced [k], but can also be pronounced [t͡s] depending on the letters that surround it, as in the start of the word *circa* ([t͡siɐkaː]). However, since letters in our study were presented in isolation, we captured each letter's dominant phoneme only, as the phoneme most likely to be associated with a single letter without surrounding context.

To calculate an RDM for phonological representations of letters' most common phonemes, we first identified each letter's dominant (most common) phoneme. We used the *epitran* library (Mortensen et al., 2018) for Python, with custom German mapping files, to perform grapheme-to-phoneme transcription of all words in SUBTLEX-DE (Brysbaert et al., 2011). Custom files were needed to clearly disambiguate the grapheme-to-phoneme transcription in cases where multiple phonemes are matched to the same character, in a manner that depends on the preceding context. This was necessary because the default epitran mapping files are designed to perform standard grapheme-to-phoneme transcription where such disambiguation is not critical.

Once transcribed, we used word frequencies to calculate the frequency with which any individual lower-case letter was transcribed to any phoneme segment in the corpus. Since we were only interested in letters in isolation (i.e., without surrounding context), we ignored cases where multiple letters combined to map to a single phoneme (e.g., *sch* mapping to ʃ). However, we did record cases in which single letters were mapped to multiple phonemes, (e.g., *x* mapping to two phonemes, [ks]). The most dominant phoneme segment for any given letter was then defined as that with the highest total frequency of transcription in the corpus.

After identifying each lower-case letter's dominant phoneme segment, we used epitran to look up corresponding vectors of 24 phonological features for each phoneme, via the *PanPhon* library (Mortensen et al., 2016). For all but one letter (*x*), the dominant phoneme segment comprised just one phoneme. For the letter *x*, the vectors for [k] and [s] were averaged. We then calculated a phonological RDM between dominant phonemes as the correlation distance between all lower-case letters' dominant phoneme vectors.

***(b) Letter Name Phonemes***

Finally, we considered that, rather than activating dominant phonemes, individual letters may have activated phonemic representations of letter *names*. For example, upon seeing the letter *ß*, participants may have activated the name for the letter, Eszett ([ɛsˈt͡sɛt]). To test this, we transcribed each of the lower-case German letters' names into IPA, based on recordings of one of the authors (RS; native German speaker) reading the letters' names aloud. We then used PanPhon to look up corresponding 24-feature phonological vectors. These vectors were then averaged for each letter, and the letter name RDM was calculated as the correlation distance between the average letter name vectors.

**Results**

All RDMs were rank-transformed, and the Bayesian multivariate model was fit as specified in the main analysis, but with these additional RDMs included. For feasibility, the time-resolved models were fit with only 4 chains rather than 8 (see **Supplementary Materials E** for a full comparison to all other models).

Results (**Figure 9**) suggested that the main analysis of posterior channels was not confounded by the control variables we considered. Rather, the partial correlations in the period of interest (**Figure 9a**) showed that Wasserstein distance still accounted for much unique variance. Time-resolved results (**9b**) showed a similar pattern.

As expected, visual size explained much of the variance in early periods, though it shared some of this variance with Jaccard distance. This is likely because the Jaccard index is constrained by the relative sizes of the letters. If two letters have very different sizes, then the maximum overlap is the case where all of the smaller letter overlaps with a part of the larger letter. To reiterate, the partial correlations suggest that, in contrast, Wasserstein distance was explaining variance that could not be accounted for by visual size.

There was no clear effect of letter frequency until later than 500 ms. Moreover, there was no clear effect of the phonological variables for the posterior electrodes.

Our priority in this analysis was to examine whether the control variables confounded our reported effect. Our findings showed that the brain-model alignment observed for Wasserstein distance could not be accounted for by the control variables. We were then interested in whether the effects of the control variables, especially of phonological effects, would have been more easily detectable had we not examined only a region of interest of posterior electrodes. Indeed, this decision was originally made to maximise sensitivity to the correlates of visual processing we were chiefly interested in. To examine effects of correlates outside of the posterior region of interest, we ran the same control analysis using RDMs calculated from all 64 scalp electrodes that we recorded from.

The scalp-wide analysis (**9b right panels**) showed very similar results to the analysis using only posterior electrodes. Notably, there was still no clear effect of the phonological variables.

**Figure 9**

*Results from the model including control variables.*

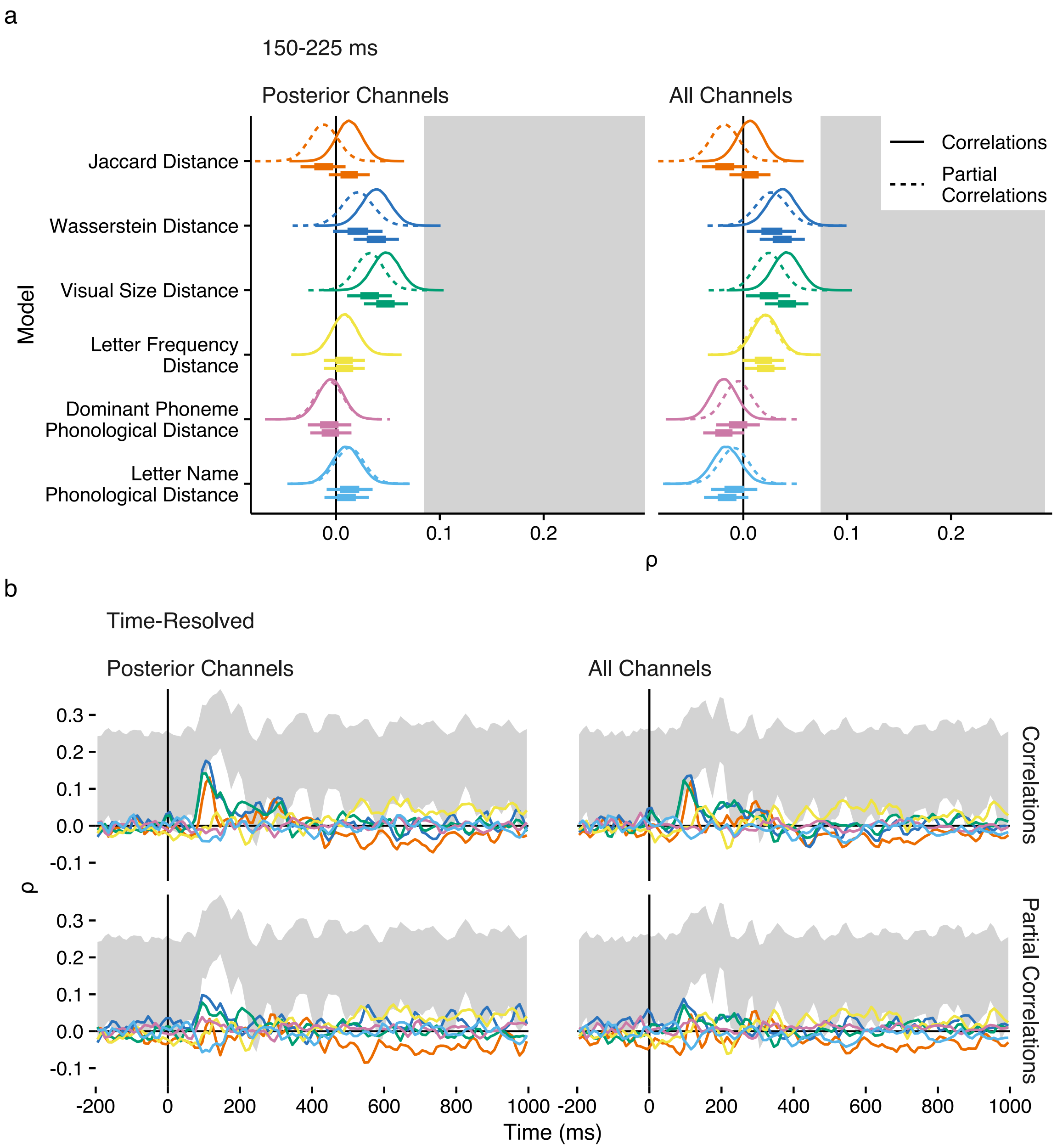


**(a)** Posterior distributions between Jaccard distance, Wasserstein distance, and all control variables' RDMs in the 150-225 ms period of interest. Boxes and whiskers below distributions show the extents of 50% and 89% HDIs. **(b)** Medians of posterior samples for all brain-model correlations and partial correlations in 10 ms windows. For both analyses, separate models were fit to neural dissimilarities calculated for (left) posterior, and (right) scalp-wide channels.

### E Bayesian Modelling Overview

We estimated the rank (Spearman's $\rho$) correlations between model and empirical neural RDMs (calculated from neural data in periods of interest or 10 ms windows). Specifically, we fit Gaussian multivariate Bayesian models. In this way, we could model the full correlation matrix between neural RDMs and either one or multiple candidate model RDMs (Kurz, 2019). This allowed us to additionally calculate a partial correlation matrix for each posterior sample. The models were fit via *brms* (Bürkner, 2017), an interface to *STAN* (STAN Development Team, 2024). As an example, the model formula for the planned analysis was specified as:

```
bf(mvbind(rank_eeg, rank_ot, rank_jacc) ~ 0) + set_rescor(rescor=TRUE)
```

In this formula, `rank_eeg` refers to the rank, within-participant neural RDMs. Similarly, `rank_ot` and `rank_jacc` refer to the rank letter similarities from the Wasserstein and Jaccard distance RDMs, which were the same for each participant. The "`~ 0`" portion of the formula specifies that the model was fit with no intercept in the average of the RDMs, as all three variables were mean-centred on zero. The `mvbind()` function was used to specify that the three variables should be modelled multivariately, and the `set_rescor()` function was used to specify that the correlations between the three variables should be estimated.

The $\sigma$ parameter (i.e., standard deviation) for all RDM variables was held constant, at the population standard deviation of all integers from 1 to 435 (i.e., 125.5734), reflecting the 435 unique dissimilarity values. This decision was made because the standard deviation for the uniform distribution of rank values is known exactly, and so we would gain no information by modelling it.

The prior for the correlation matrix was specified as a Lewandowski-Kurowicka-Joe (LKJ) distribution of $\eta$=1.5, resulting in a distribution biased somewhat towards 0, and away from -1 or 1, with around 80% of the probability mass in the marginal distributions between -.6 and .6. This distribution was chosen to reflect our expectation that the correlation would probably be quite small, informed by the high degree of noise inherent to EEG, and by the magnitude of brain-model correlations in previous RSA analyses of EEG data (e.g., He et al., 2022; Salmela et al., 2018), while still allowing for somewhat higher correlations between model RDMs (e.g., between Wasserstein and Jaccard distance). We conducted a sensitivity analysis (**Supplementary Materials F**) which additionally showed that the observed pattern of results was robust to even priors biased very strongly towards correlations of zero or one.

The `adapt_delta` parameter was set to .99 for all models.

In **Table 1**, we summarise all the parameters selected and variables modelled in the Bayesian RSA models we reported.

| Manuscript Section and Model Summary | chains | Iterations | |
|---|---|---|---|
| | | warmup | sampling |
| Planned Analysis<br>Modelled the correlation matrix between neural RDMs, Jaccard distance, and Wasserstein distance in the 150-225 ms period of interest. | 8 | 6,000 | 18,000 |
| Time-Course Analysis<br>Time-resolved models, each modelling the correlation matrix between neural RDMs, Jaccard distance, and Wasserstein distance in 10 ms windows. | 8 | 5,000 | 5,000 |
| Analysis of Geometric Invariance<br>Individual models of the correlation between neural RDMs and a given variant of Jaccard or Wasserstein distance. Individual models were fit to the 150-225 ms period of interest, an additional 80-130 ms period, and to each 10 ms window in the neural data. | 8 | 5,000 | 5,000 |
| Comparison to ANNs (periods of interest)<br>Individual models of the correlation between neural RDMs and a features from a given module of each variant of ResNet-50 and CORnet-Z. Individual models were fit to the 150-225 ms period of interest and the 80-130 ms period. | 8 | 5,000 | 5,000 |
| Comparison to ANNs (time-resolved)<br>As described above, but fit individually to each of the 10 ms windows. For feasibility, the number of chains was reduced to 4 per model. | 4 | 5,000 | 5,000 |
| Supplementary Materials D (period of interest)<br>Modelled the correlation matrix between neural RDMs, Jaccard distance, and Wasserstein distance, and 5 control RDMs in the 150-225 ms period of interest. | 8 | 6,000 | 18,000 |
| Supplementary Materials D (time-resolved)<br>Modelled the correlation matrix between neural RDMs, Jaccard distance, and Wasserstein distance, and 5 control RDMs in 10 ms windows. For feasibility, the number of chains was reduced to 4 per model. | 4 | 5,000 | 5,000 |

**Table 1**
*All parameters that varied between the fitted models. Columns chains, warmup, and sampling correspond to the STAN parameters of the same name.*

### F Sensitivity Analysis

As outlined in **Supplementary Materials E**, we specified a Lewandowski-Kurowicka-Joe (LKJ) prior distribution of $\eta$=1.5 for the correlation matrix. Values of $\eta$ below 1 bias marginal correlations symmetrically towards extreme values of -1 and 1, whereas $\eta$ values above one bias correlations symmetrically towards zero (**Figure 10a**). Our chosen value of $\eta$=1.5 biased the estimated correlations slightly towards zero, reflecting our expectation that the correlations with the neural data would be quite weak, while accommodating a medium correlation between Wasserstein and Jaccard distance.

**Figure 10**
*Sensitivity analysis.*

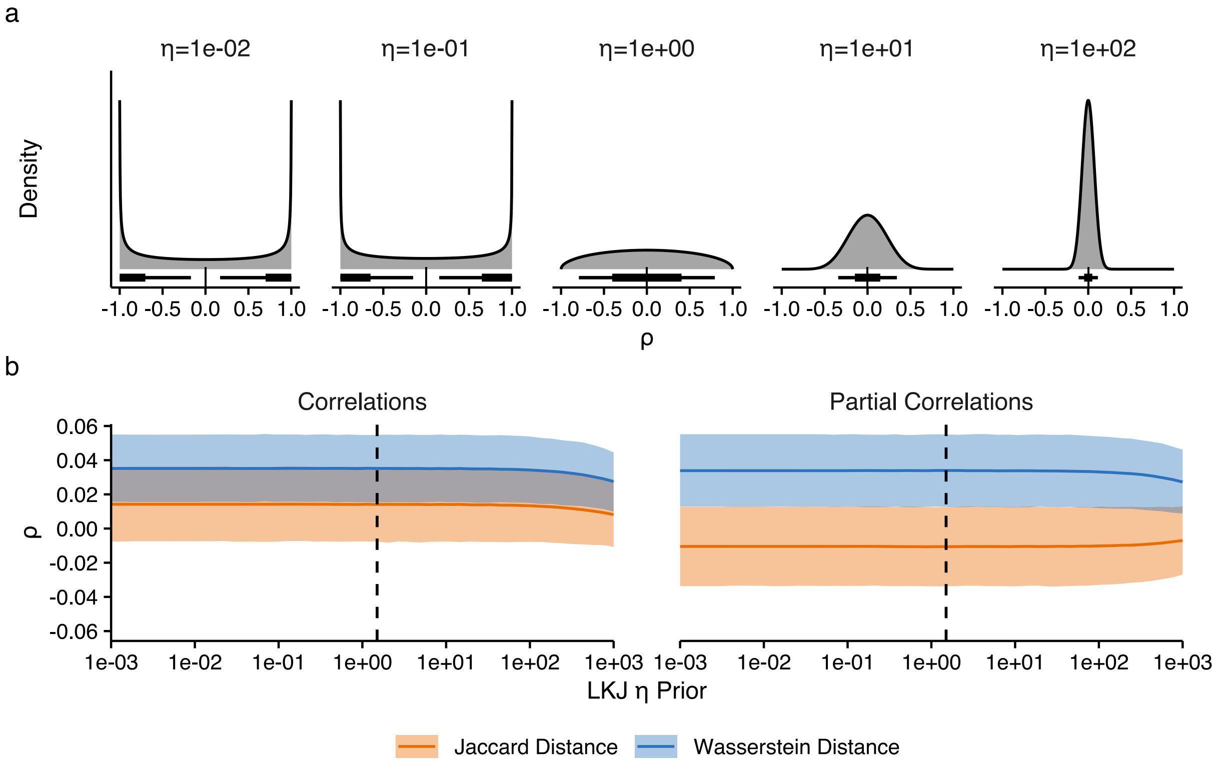


**(a)** Illustration of the effect of varying the $\eta$ parameter on the shape of a marginal LKJ distribution, for a correlation matrix with three variables. Vertical lines below distributions reflect the median (which is always zero), while boxes and whiskers reflect the extents of 50% and 89% HDIs respectively. **(b)** Effect of varying the $\eta$ parameter for the prior on the posterior estimates for correlations between neural RDMs and Jaccard and Wasserstein distance. Central lines depict medians, while shaded regions depict 89% HDIs. The dashed line depicts the location of $\eta$=1.5 used in our analysis.

To examine how sensitive our posterior estimates were to our choice of prior, we conducted a sensitivity analysis. Here, we fit models exactly as described for the planned analysis, but varying the LKJ prior's $\eta$ value. Specifically, we fit 50 models with $\eta$ values spaced evenly on a $log_{10}$ scale from 1e-3 to 1e3. Results (**Figure 10b**) showed that the estimates were very stable, with only extremely biased priors greatly influencing the results.

### G Details on ANN Model Training and Feature Extraction

We examined alignment between neural RDMs and those calculated from layers of two artificial neural networks (ANNs) - ResNet-50 1.5 and CORnet-Z. Three variants of each model were trained in *pytorch* and *torchvision*:

(1) Variant trained on images of lower-case Arial Light German letters.

(2) Variant trained on images from ImageNet (i.e., simply importing the pretrained ResNet-50 weights available in `torchvision.models.ResNet50_Weights.IMAGENET1K_V2`, and the pretrained CORnet-Z weights available at https://s3.amazonaws.com/cornet-models/cornet_z-5c427c9c.pth).

(3) Variant trained on images from ImageNet (i.e., initialised with the pretrained weights mentioned in (2)), and then concurrently, with equal probability of selection, on images from ImageNet and images of Arial Light letters.

#### Training

All input images were presented as raster images at a resolution of 224×224 pixels, with a datatype of `torch.float32`. All images in the training set had red, green, and blue channels, but images of letters were identical in all three channels. For comparability with the models pretrained on ImageNet, images were normalised using the mean and standard deviation of the colour channels of images in the ImageNet dataset.

When images of letters were presented in the training set, they were presented at a random font size between a maximum of 15 and 250 pixels (varied by integer, and no letters used the maximum space available). Letters were also presented with a random rotation between -15 and 15 degrees. Letters were presented in random locations in the images, though such that no non-zero pixels were outside of the image's 224×224 bounds. When images from ImageNet were presented in the training set, they were cropped to fit the input dimensions at random locations. There was additionally a 50% chance that an image from ImageNet might be flipped horizontally.

ResNet-50 models were trained using an Adam optimiser, with a learning rate of .001.

ResNet-50 was trained in 15 epochs, with a batch size of 64. CORnet-Z models were trained using a stochastic gradient descent optimiser, with an initial learning rate of .01, a momentum of .9, and a weight decay of .0001. CORnet-Z was trained in 15 epochs, with a batch size of 256. The step size for CORnet-Z was decreased by 10 times every 10 epochs. All models were trained using a cross-entropy loss function.

**Feature Extraction and RDM Calculation**

We created images of all lower-case German letters in Arial light font at a maximum size of 250 pixels. The letters were presented at the centre of images with 224×224 input dimensions, with identical red, green, and blue channels, and a data type of `torch.float32`. As in training, images were normalised using the mean and standard deviation of the colour channels of images in the ImageNet dataset.

We first used these letter images to check accuracy in model predictions output by the four models trained to recognise letters. Three of four models showed 100% (top-1) accuracy in letter identities predicted from the model output layers. The ResNet-50 model trained on letters only, which made one incorrect prediction of *n* for the letter image of *m*. Nevertheless, *m* and *n* showed very similar activation, and *m* was the category predicted with the next greatest activation after *n*.

We next constructed RDMs from features extracted from ANNs via hooks attached to intermediate modules of the models, as implemented in the `torchextractor` Python library (https://github.com/antoinebrl/torchextractor). For ResNet-50, activations were extracted from the blocks that make up each of the four layers: layer 1 / conv_2 (3 blocks), layer 2 / conv_3 (4 blocks), layer 3 / conv_4 (6 blocks), and layer 4 / conv_5 (3 blocks). This resulted in activations from a total of 14 blocks. For the shallower architecture of the CORnet-Z models, activations were extracted from the model's four layers: V1, V2, V4, and IT. Finally, raw RDMs for a given block or layer were calculated as Pearson correlation distances between the activations elicited by pairs of letters being compared. These RDMs were then rank-transformed for the RSA.
